# Solid state anaerobic digestion of mixed organic waste: the synergistic effect of food waste addition on the destruction of paper and cardboard

**DOI:** 10.1101/564203

**Authors:** Nigel G. H. Guilford, HyunWoo Peter Lee, Kärt Kanger, Torsten Meyer, Elizabeth A. Edwards

## Abstract

Full-scale anaerobic digestion processes for organic solid waste are common in Europe, but generally unaffordable in Canada and the United States because of inadequate regulations to restrict cheaper forms of disposal, particularly landfill. We investigated the viability of solid-state anaerobic digestion (SS-AD) as an alternative that reduces the costs of waste pretreatment and subsequent wastewater treatment. A laboratory SS-AD digester, comprising six 10L leach beds and an upflow anaerobic sludge blanket reactor treating the leachate, was operated continuously for 88 weeks, with a mass balance of 101±2%. The feed was a mixture of cardboard, boxboard, newsprint, and fine paper, and varying amounts of food waste (from 0% to 29% on a COD basis). No process upset or instability was observed. The addition of food waste showed a synergistic effect, raising CH_4_ production from the fibre mixture from 52.7 L.kg^-1^COD fibre_added_ to 152 L.kg^-1^COD fibre_added_, an increase of 190%. Substrate COD destruction efficiency reached 65% and a methane yield of 225 L.kg^-1^ COD_added_ was achieved at 29% food waste on a COD basis, and a solids retention time of 42 days. This performance was similar to that of a completely stirred tank reactor digesting similar wastes, but with much lower energy input.

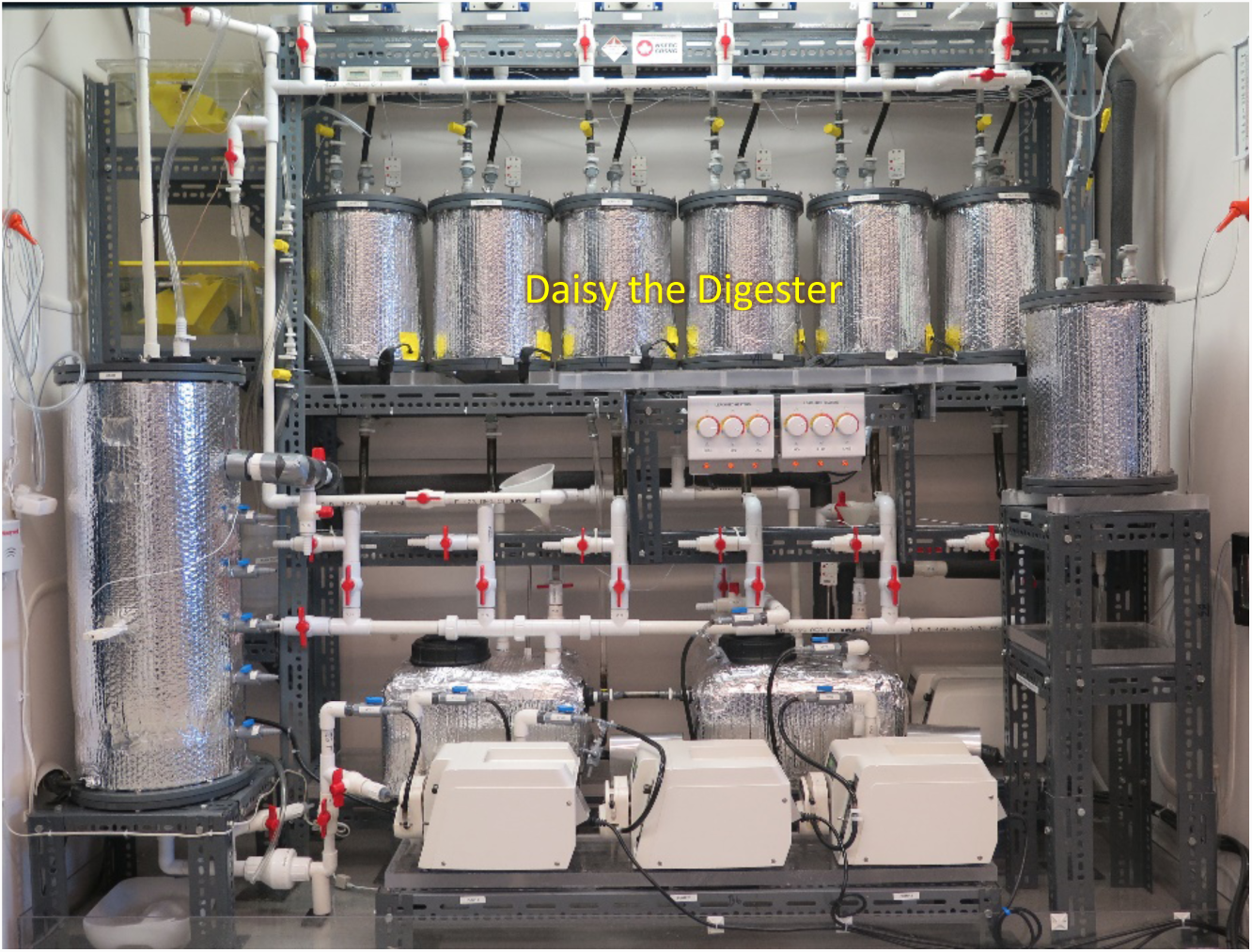

## 1. INTRODUCTION

Anaerobic digestion (AD) is a well-established method for the treatment of organic solid waste and the production of renewable energy. In a comparative study, Hodge et al. (2016) concluded that, among composting, landfilling, combustion with energy recovery and AD, a combination of AD and landfill was the leading alternative in terms of lowering global warming potential. Solid organic waste is heterogeneous, variable and complex. Consequently, conventional anaerobic digesters (De Baere & Mattheeuws, 2013; Guilford, 2009) must be preceded by extensive pretreatment, such as sorting, size reduction, contaminant removal, and water addition, to render the feedstock suitable for processing, all of which add significantly to the cost.

The Landfill Directive of the European Union (1999) restricts the disposal of organic wastes, and has driven the widespread adoption of more expensive waste processing technologies, particularly anaerobic digestion. Canada and the United States lack similar, overarching, regulations; consequently, anaerobic digestion is common in Europe and relatively rare in North America. As a direct result, in Canada, most organic waste (about 10 million tonnes per year (Government of Canada, 2015) consisting of about 38% food waste 62% paper and cardboard (City of Ottawa, 2007; Government of Ontario, 2004) is still disposed of in landfills which, in the aggregate, generate about 20 million tonnes per year of greenhouse gases (CO_2_ eq.) (Environment Canada, 2017); furthermore, a source of renewable energy is largely forgone. Satchwell et al. (2018) note the advantages of solid-state anaerobic digestion (SS-AD), including less pretreatment and reduced wastewater treatment, but identify numerous scientific, operational, and policy challenges limiting its wider adoption in the United States and Canada.

In an attempt to circumvent the lack of strong regulations, or incentives, in North America (Guilford, 2017), a new approach to AD was developed and patented to treat all forms of solid organic waste from residential and commercial sources (Forrestal et al., 2006a; Forrestal et al., 2006b). The underlying principle, derived from bioreactor landfill practices, is to accommodate the properties of the waste as-received as far as possible. The process employs SS-AD; the waste remains stationary, and the leachate generated by its degradation is recirculated through the waste. Biogas is recovered and put to beneficial use, and the digestate remaining is aerobically cured and turned into compost. It is designed to accommodate the complex properties of solid waste with minimal pretreatment, with the ultimate goal of being cost-competitive with landfill; initial estimates suggested that this can be achieved (Guilford, 2009). In exchange for simplicity of design, some trade-offs were expected. For example, it was assumed that a lower substrate destruction efficiency would be achieved, with a longer solids retention time (SRT), and a larger physical footprint required (compared to, for example, a CSTR), but that capital and operating costs would be lower.

The AD technologies commonly applied to solid waste employ various configurations and operating conditions (De Baere & Mattheeuws, 2013). The majority are single stage, and either plug flow or CSTRs, operating at 38°C or 55°C. They digest primarily food waste (FW) and the organic fraction of municipal solid waste (OFMSW), plus some leaf and yard waste in some cases. Consequently, most of the research on the AD of organic solid waste uses FW or OFMSW as the substrate. Zhang et al. (2012) measured the digestibility of OFMSW in a CSTR, giving 62% substrate destruction as volatile solids (VS) and yielding 304 L CH_4_/kg VS_added_; Browne et al. (2013a; 2013b; 2014) tested a two-stage digester comprising leach beds (LBs) and upflow anaerobic sludge blanket (UASB), giving 75% substrate destruction as VS digesting OFMSW and yielding 384 L CH_4_/kg VS_added_, but experienced serious problems with hydraulic conductivity, ammonia inhibition, and volatile fatty acid (VFA) inhibition. Though lignocellulosic fibers make up a high proportion of solid organic waste, only a very few studies have examined the digestibility of these substrates (Di Maria et al., 2017; Eleazer et al., 1997; Pommier et al., 2010; Yuan et al., 2012; Yuan et al., 2014). These previous experiments are summarized in Results and Discussion, Section 3.8.

To evaluate the effectiveness of SS-AD, we designed and built Daisy the Digester, a lab-scale version of the new SS-AD digester design (Guilford, 2009), a hybrid system combining the robustness and simplicity of a landfill bioreactor with the benefits of multi-stage digestion. Daisy comprises six sequentially-fed leach beds, an upflow anaerobic sludge blanket (UASB) and two tanks, plus ancillary components and a control system. We also designed a feed stream composed of a mixture of cardboard (CB), boxboard (BB), newsprint (NP) and fine paper (FP), collectively representing the fibre fraction (FB), plus varying amounts of food waste, to simulate the range of composition of industrial, commercial and institutional waste (IC&I) (Government of Canada, 2010). In order to maintain permeability, shredded ash wood was added as a bulking agent (BA). The objectives of the research were to measure process stability and digester performance (methane yield and substrate destruction efficiency) in response to variations in the proportion of food waste added, for comparison with more conventional CSTR-type wet digesters processing similar wastes. As a result of extensive careful and frequent monitoring of the system, the mass balance over the entire 88-week experiment was nearly perfectly conserved, revealing a remarkably strong effect of food waste on the extent of digestion of the fiber fraction.

## 2. MATERIALS AND METHODS

### 2.1 Daisy the Digester – Design

The design is derived from bioreactor landfill practice, in which leachate recirculation accelerates the decomposition of unsorted solid waste (Guilford, 2009). Daisy comprises six 10L leach beds (8.5L working volume), a UASB (27.5L working volume), a UASB feed tank (Tank 1) and a leach bed feed tank (Tank 2) (17.5L working volume each), three peristaltic pumps (P1, P2, and P3) and two wet-tip gas meters (GM1 and GM2) to measure biogas volumes produced (Fig. 1). The tanks and the UASB are heated automatically with self-regulating heat tape; each has a programmable thermostatic controller; the LBs have manually controlled heaters. The frequency and volume of leachate delivered to the leach beds is controlled automatically with a programmable logic controller (PLC); leachate is recirculated from Tank 2, via P3, through the upper manifold, and back to Tank 2. Periodically, the automatic valves, controlled by the PLC, open in sequence and redirect the flow to each leach bed in turn. The cycle repeats continuously. Leachate drains from each leach bed to a common manifold and into Tank 1; it is then transferred by P1, via a clarifier within the tank, to the UASB. Effluent from the UASB is discharged to Tank 2. The hydraulic balance between Tank 1 and Tank 2 is maintained by P2 and an overflow pipe connecting the two. Hydraulically, Daisy is a closed system. Biogas from the UASB is discharged through GM1, and the aggregated biogas from six LBs and two tanks is discharged through GM2. Daisy runs at a slight positive pressure of 1.2 kPa, generated by 12 cm water column in gas meters. Fig 1B shows 9 liquid sampling valves; V2 (UASB feed) was used for all the samples reported here; the balance were used in a companion study (Lee, 2018) intended for publication at a later date. The design basis, construction, and operation of Daisy are described in much greater detail elsewhere (Guilford, 2017).

**Fig. 1:**
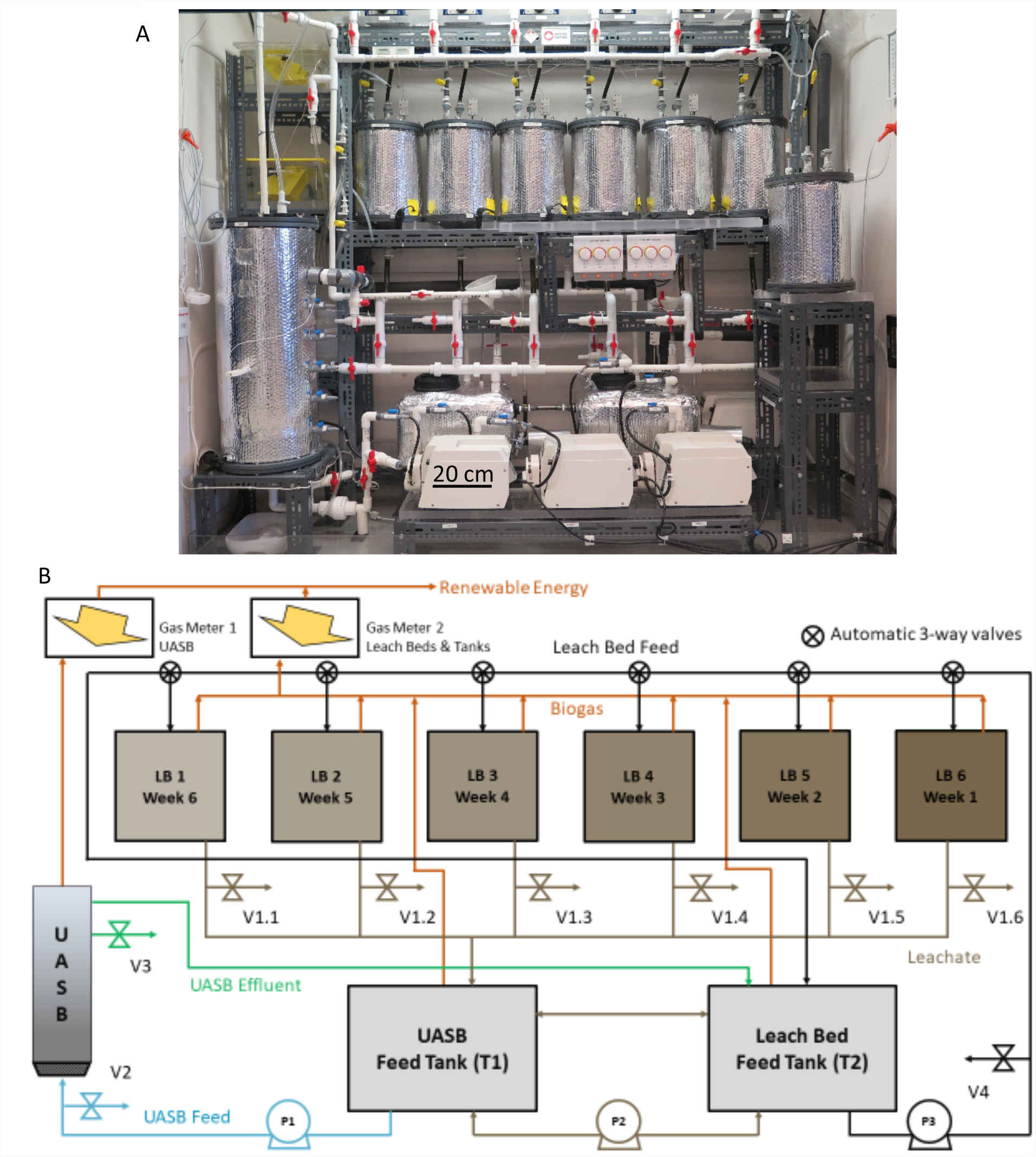
Daisy the Digester. A. Photograph; B. Schematic process flow diagram; 6 LBs fed sequentially at 1 wk. intervals; one UASB; 2 leachate tanks – T1 to feed UASB, and T2 to feed LBs; 3 peristaltic pumps – P3 feeding LBs, P1 feeding UASB, P2 balancing T1 and T2; two wet-tip gas meters, 6 automatic 3-way valves and 9 sampling valves.

### 2.2 Operational set-up of Daisy

Daisy operated as a sequentially-fed batch reactor with a fresh LB of waste (containing 1.2 to 1.7 kg of substrate as COD) added once per week, replacing a 6-week old leach bed that was removed for analysis. As depicted in Fig. 1B, LB 1 is due for replacement after an SRT of 42 days. The UASB and Tank 1 were set at 37°C; Tank 2 was set at 39°C (this was to provide additional heat to the LBs, before they were equipped with manual heaters at week 25). The leachate recirculation rate remained constant throughout at 565 mL per LB, every 30 minutes. This value was derived from Murto et al. (2013) who reported a flowrate of 16.5 L.min^-1^ in a 5.2 m^3^ leach bed containing 3.4 t of waste plus 2.6 m^3^ of water, or 2.8 t of a mixture of waste and bulking agent plus 2.6 m^3^ of water. The UASB was fed at 125 mL.min^-1^ giving a hydraulic retention time (HRT) of 3.6h and an upflow velocity (V_up_) of 0.5 m.h^-1^. The peristaltic pumps were calibrated with and without a 1.3m head, with new tubing and with worn tubing (Norprene A-60-G); the calibration remained unchanged. The wet tip meters (GM1 and GM2), supplied by Archaea Press, were calibrated using a continuous water-displacement method (Guilford, 2017); biogas production (100mL/tip) was recorded in the datalogger every 5 minutes and also every hour. Temperature was recorded every 15 minutes (six LBs, two tanks, GM2 and the UASB). Biogas volumes were corrected to STP using the recorded temperature inside GM2, and the barometric pressure was recorded by a weather station located on the roof of the building.

### 2.3 Feedstock and digestate – preparation, sampling and analysis

All components of the feedstock were recovered from residential waste recycling programs and prepared as follows. The CB and BB were coarsely shredded (<3cm × 4cm); the FP and NP were shredded in an office shredder (< 5cm × 0.5 cm); the BA, consisting of prepared ash wood, was supplied in 6 batches (BA#1 to BA#6). BA#5 was processed through a chipper; the other five were shredded in a Roto-Chopper and screened to <5cm; all were stored in bulk. The FW was recovered from a residential green bin program in the Region of Durham, Ontario (which prohibits sanitary products and non-compostable plastic). It was presorted to remove plastic and larger junk, shredded to <10cm in a shear shredder, and stored in sealed plastic bags (∼ 1.5kg ea.) at −20°C.

The FW was thawed as needed, and hand-sorted to remove bones, inorganic matter, and smaller foreign objects; it was either fed directly to Daisy (weeks 12 to 76), or first pulped in a blender with an equal quantity of water (weeks 1 to 11 and 77 to 88). The fibres (FB) and BA were weighed, and mixed dry, in a 20L bucket using large stainless-steel spoons. Water was added to saturate the fibres (between 3.8L and 3.2L depending on the amount of food waste added); the FW was added last and thoroughly mixed in using the same method.

To measure the digestibility of individual components of the feed, at different levels of FW addition under actual digester conditions, stainless steel tea balls or ‘coupons’ (Fig. S1), were filled with samples of a single fibre (between 1 and 4g), and inserted into the waste. Thus, at any given moment, four of the LBs each contained six 2.5 cm tea balls comprising two triplicate sets; for example, three of CB and three of NP, or three of BB and three of FP. The other two LBs each contained a single 5 cm tea ball containing a sample of BA (7 to 11g); the larger size was necessitated by the morphology of the BA.

Each week fresh waste (feed) was placed into a LB, tamped down by hand, the head space measured, the lid installed, and the assembly flushed, leak tested, and pressurized to 50 cm water column (WC), with argon, before installation in Daisy. Quick-disconnect fittings with shut-off valves enabled rapid LB removal and replacement without ingress of air. At the end of each digestion period (typically 6 weeks), a LB was removed and replaced with a LB of fresh waste. After removal, each LB was drained for 24h, and the recovered leachate returned to Tank 1 (through a valved port to preserve gas pressure). The headspace was re-measured and the settlement noted. The coupons were removed and weighed and their TS/VS determined using standard methods; 50 mL samples of the digestate (DG) were taken from 13, 20, 25 and 28 cm from the top of the LB, for determination of TS/VS; a separate sample (also taken at 25 cm) was analyzed for COD. A 300g bulk sample of DG was retained and frozen at −20°C. A detailed record of the input and output for every LB was maintained. An example is shown in Table S1.

### 2.4 Experimental Design

The 88-week experiment was divided into 12 periods, each representing a different set of operating conditions (Table 1). After initial start-up, which took 5 weeks, the impact of specific process changes was investigated. In Period 1 (weeks 6 to 15) consistent operation was established. The solids retention time (SRT) was always set at 42d except during Period 2 (weeks 16 to 24) which briefly explored an increase in SRT to 49d (7 weeks) by omitting LB replacement every 6th week; in Period 3 Daisy was returned to 42d (6 week) SRT; COD_FW_ addition was 17.2% throughout Periods 1, 2 and 3. In Periods 4a, 4b, and 4c, COD_FW_ addition was reduced to 12.9%, 7.9% and 0% respectively. In Period 5, COD_FW_ addition was returned to 17.2% in a single step; a change to a new batch of bulking agent at week 58 caused a decline in performance which took 15 weeks to resolve. In Period 6a COD_FW_ addition was increased to 21.2% and in Period 6b to 29.3%.

**Table 1.** Operating conditions, substrate destruction efficiency as VS and COD, and CH_4_ yield, all by Period

| Operating Conditions |  |  |  |  |  |  |  |  | % Substrate Destruction |  | CH<sub>4</sub> production |  |  |  |
| --- | --- | --- | --- | --- | --- | --- | --- | --- | --- | --- | --- | --- | --- | --- |
| Period | Weeks | BA Batch | gCOD<sub>added</sub>/LB |  |  | <sup>b</sup>COD<sub>FW</sub> % | <sup>c</sup>C:N ratio | SRT days | VS | COD | L.wk<sup>-1</sup> | n | L.kg<sup>-1</sup> VS<sub>added</sub> | L.kg<sup>-1</sup> COD<sub>added</sub> |
|  |  |  | BA | <sup>a</sup>FB | FW |  |  |  |  |  |  |  |  |  |
| 1 | 6 - 15 | 1 | 645 | 1200 | 250 | 17.2 | 74 | 42 | 57.3 | 53.7 | 247±7 | 10 | 219 | 169 |
| 2<sup>d</sup> | 16 - 24 | 2 | 645 | 1200 | 250 | 17.2 | 74 | 49 | 58.2 | 53.5 | 197±26 | 9 | 223 | 172 |
| 3 | 26 - 31 | 2 & 3 | 645 | 1200 | 250 | 17.2 | 74 | 42 | 59.3 | 54.2 | 278±18 | 5 | 239 | 185 |
| 4a | 32 - 37 | 4 | 645 | 1200 | 178 | 12.9 | 93 | 42 | 43.2 | 38.9 | 198±5 | 5 | 184 | 143 |
| 4b | 38 - 44 | 4 | 645 | 1200 | 104 | 7.9 | 130 | 42 | 33.2 | 31.8 | 138±3 | 4 | 135 | 105 |
| 4c | 44 - 49 | 4 | 645 | 1200 | 0 | 0.0 | 350 | 42 | 20.0 | 18.6 | 63±4 | 4 | 67 | 52.7 |
| 5a | 50 - 57 | 4 | 645 | 1200 | 250 | 17.2 | 74 | 42 | 53.1 | 51.7 | 217±21 | 6 | 210 | 162 |
| 5b | 58 - 63 | 5 | 645 | 1200 | 250 | 17.2 | 74 | 42 | 45.9 | 43.8 | 251±26 | 6 | 217 | 168 |
| 5c | 64 - 70 | 4 | 645 | 1200 | 250 | 17.2 | 74 | 42 | 48.7 | 43.9 | 218±16 | 6 | 200 | 154 |
| 5d | 71 - 74 | 6 | 645 | 1200 | 250 | 17.2 | 74 | 42 | 56.3 | 53.0 | 245±19 | 5 | 219 | 169 |
| 6a | 74 - 80 | 6 | 645 | 1200 | 333 | 21.7 | 62 | 42 | 63.5 | 56.0 | 297±12 | 6 | 246 | 189 |
| 6b | 81 - 88 | 6 | 645 | 1200 | 500 | 29.3 | 48 | 42 | 69.4 | 65.3 | 384±8 | 6 | 296 | 225 |

<sup>a</sup>gCOD<sub>FB</sub> comprises CB = 482, BB = 380, NP = 142 and FP = 196; <sup>b</sup>Calculated as percent of FB + FW (without BA). <sup>c</sup>Calculated as a weighted avg. from elemental analysis (Table S2) <sup>d</sup>Seven week SRT achieved by twice skipping a LB replacement at the seventh week, over a total of 9 weeks; substrate destruction efficiency unchanged; 7 weeks of methane production spread over 9 weeks to give an effective yield = 197\*9/7 = 253 L.wk<sup>-1</sup>; cf Period 1.

### 2.5 Sampling and Analytical Methods

The sampling, analytical and data management methods used are summarized below; a more detailed description is provided in the Supporting Information (SI). The elemental composition of each individual component of the feedstock was determined using a Thermoflash 2000 CHN analyzer; TKN was also measured to achieve greater precision for nitrogen. Total solids (TS), volatile solids (VS) and chemical oxygen demand (COD) were measured using standard methods (APHA, 1992); VS was used for comparison with published results, COD measurements formed the basis of the mass balance calculation. The COD content of all feedstocks was also calculated from their elemental composition, for comparison to the measured values.

Four leachate samples were withdrawn from valve V2 (Fig. 1B) four times per week. One (15 mL) was analyzed for TS, VS, and COD using standard methods (APHA 92); the second (50 mL) to determine the pH and alkalinity ratio, the third (2×10 mL) was prepared and stored for subsequent microbial analysis and the fourth (10 mL) was filtered using 0.22µm nylon syringe filter and stored at −20°C for subsequent ion chromatography (IC) analysis for VFAs and sulphate. Sampling for VFA and microbial analysis began at week 35. Approximately every two weeks, samples of biogas (200 µL) were extracted through septa, installed in the infeed lines to GM1 and GM2, and analyzed for CH_4_ and CO_2_ using a gas chromatograph with a thermal conductivity detector (GC-TCD). Temperatures and biogas volumes were recorded in the data logger and downloaded daily. A daily activity log was maintained to record all inputs and outputs (time, type and volume), system adjustments, operating anomalies, and corrective measures.

### 2.6 Calculations

The stoichiometric formula of the substrates *C*_*n*_*H*_*a*_*O*_*b*_*N*_*c*_, and the stoichiometry of digestion, were calculated from elemental analysis of the weighted average composition of the substrates fed to Daisy (weeks 6 to 88 inclusive) using Equations (1-4) (Rittmann & McCarty, 2001).

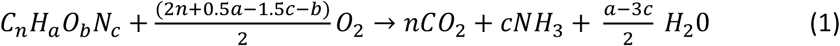

The COD of each substrate was calculated from:

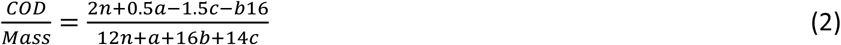

where:

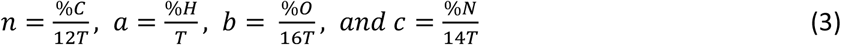

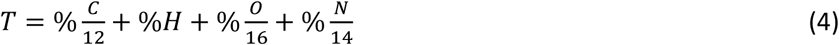

The C:N ratio was calculated from:

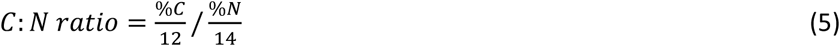

The mass balance (on a COD basis) was calculated in two ways, Method A: (methane + new biomass)/ COD_destroyed_, and Method B: COD_products_/COD_substrates_:

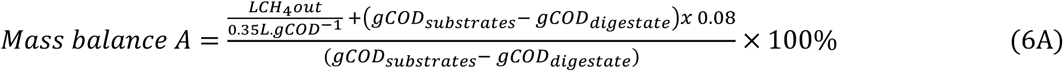

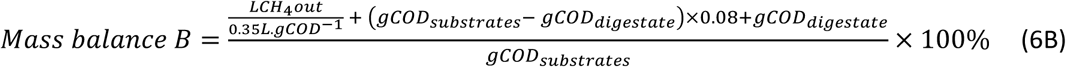

Substrate destruction efficiency was calculated from:

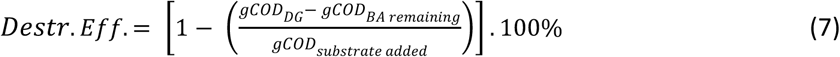

where:

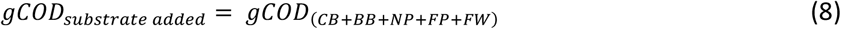

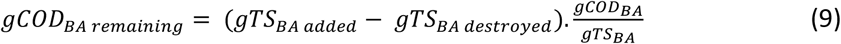

Specific methane yield, by period, was calculated from:

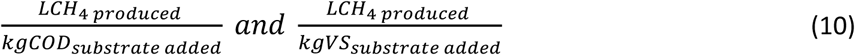

Synergistic biogas was calculated from:

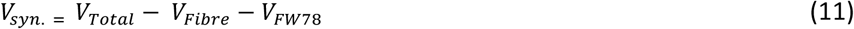

Where: V_syn._ is the synergistic (or unaccounted for) methane generated from fibre, V_total_ is the measured total methane produced; V_fibre_ is the measured methane produced at 0% COD_FW_, and V_FW78_ is the calculated maximum volume of methane generated from the added FW alone, from COD_FWconverted_, assuming 78±1% COD_FW_ conversion, a value obtained from our biochemical methane potential (BMP) tests (Guilford, 2017) in agreement with the literature (Eleazer et al., 1997).

## 3. RESULTS AND DISCUSSION

The results are described and discussed from seven perspectives; 1) analytical results; 2) mass balance; long-term performance and stability; 4) the effect of food waste(FW) addition on the digestibility of lignocellulosic fibres, and on performance; 5) the relative digestibility of the fibres – CB, BB, NP, FP and BA – from coupon data; 6) the unexpected effect of a change in bulking agent; and 7) the effect of SRT on performance.

### 3.1 Analytical results for feedstocks, digestate and biogas

The elemental composition and ash content (and thus VS), of each the substrates and digestates, were measured and averaged; the stoichiometric formula of each substrate was calculated (Table S2); the stoichiometric formula of the 83-week weighted average feed to Daisy was also calculated as C_90_H_155_O_67_N; The COD content of each of the substrates was calculated from Equations (1), (2), (3) and (4), and compared to the measured values (Table S2). The measured and calculated values of COD content corresponded well; unsurprisingly, the greatest discrepancy was for FW, the most variable of the substrates. The TS, VS, and COD of the digestate from all 87 LBs was measured (Table S3). The average methane content of the biogas was 58.5±3.7% from GM1 (the UASB), and 51.7±3.6% from GM2 (balance of the system); the weighted average was 52.4% (Table S4). The methane content was also calculated, from digestion stoichiometry, as 52.5% (using equations shown in Fig. S3). The measured weekly volume of biogas and of CH_4_, were corrected to STP (273K and 100kPa) (Table S5).

### 3.2 Mass Balance on a COD Basis

Daisy was fed a total of approximately 97 kg of TS and 125 kg of COD over 83 weeks, and produced approximately 20,000 L of CH_4_ at STP. The mass balance (COD basis) of the entire system was calculated for each period two different ways using equations 6A and 6B, from week 6 to week 88 inclusive (Table S6). The cumulative mass balance for all 83 weeks was 101±2% (Method A) and 100±2% (Method B); these results thus validate the sampling and analytical methods used, and create a sound foundation upon which to assess Daisy’s performance. The mass balance does show a little variability when considered by individual Period, particularly using Method B (Table S6); the reasons are discussed in the description of Table S6 on page 4 of the SI.

### 3.3 Long term operation

For each of the 12 operating periods, the feedstock composition, operating conditions, and Daisy’s performance measured as substrate destruction efficiency (Equations 7, 8 and 9) and methane yield (Equation 10), are shown in Table 1. The input data to all calculations are derived from Tables S1-S5. The destruction efficiency of BA over 6 weeks averaged about 7%, irrespective of food waste addition (Table S7). Since BA is to be reused at commercial scale, and would artificially depress measurements of performance, it was excluded from the calculation of substrate destruction efficiency as shown in equations (7), (8), and (9).

The SRT remained at 42 days (*i.e.*, 6 weeks), except during Period 2 (which lasted only 8 weeks) when it was 49 days. Food waste addition, expressed as a percent of total COD added, varied from 17.2% down to 0% then back up to 29.3%. The C:N ratio, calculated from Equation 5 (Table 1), ranged between 48:1 and 350:1, depending on FW addition, and was always well above the generally accepted stability threshold of 20:1 (Igoni et al., 2008; Wu et al., 2010; Yadvika, 2004). During Period 5, the BA was changed to a different batch which, unexpectedly, reduced digester performance.

Fig. 2 shows the entirety of the experimental period, week by week, expressed as: a) substrate destruction efficiency and specific CH_4_ yield; b) alkalinity ratio and pH; c) concentration of total VFAs as COD in mg.L^-1^; d) recirculating concentration of inorganic salts in g.L^-1^; and e) recirculating concentration of SO4^2-^ in mg.L^-1^. Methane yield ranged from a low of 52.7 L.kg^-1^COD_added_ to a high of 225.4 L.kg^-^ 1COD_added_ and the corresponding COD destruction efficiency from 18.6% to 65.3% (Fig. 2A). Despite wide variations in methane yield and substrate destruction, Daisy’s operation remained stable throughout. In particular, the alkalinity ratio (weekly average) remained below 0.52 (against a target of ≤0.4) and the pH between 6.7 and 7.3, with a brief excursion to 7.6 (Fig. 2B). VFA’s and sulphate were measured four times per week, beginning at week 35. The first VFA measurement, taken 6h after installation of a LB of fresh waste, showed a sharp spike (except at zero FW); the second and third, taken 1d and 3d later, showed sharp declines (Fig. 2C). At no time was there any indication of a build-up of VFAs, hence the stability of pH and alkalinity ratio. It was discovered, by about week 20, that there was no accumulation of leachate within Daisy, and thus no free wastewater being produced. Measurement of the TS content of the digestate revealed that the same quantity of water was being removed in the digestate as was being added in the feed. By measuring the VS content of the leachate it was possible to determine the fate of the inorganic salts; their concentration within Daisy (Fig. 2D) varied with FW addition, falling from 3.5 g.L^-1^ in Periods 1, 2 and 3 to 2.0 g.L^-1^ in Period 4c, finally rising to 3.3 g.L^-1^during periods 5 and 6. The amount of inorganic matter removed in the digestate (∼ 160g per LB) approximately equaled the amount added in the feed. Up to week 50, the concentration of sulphate remained close to 50 mg.L^-1^ but then began to rise as the proportion of FW increased, ultimately reaching 125 mg.L^-1^ (Fig. 2E).

**Fig. 2:**
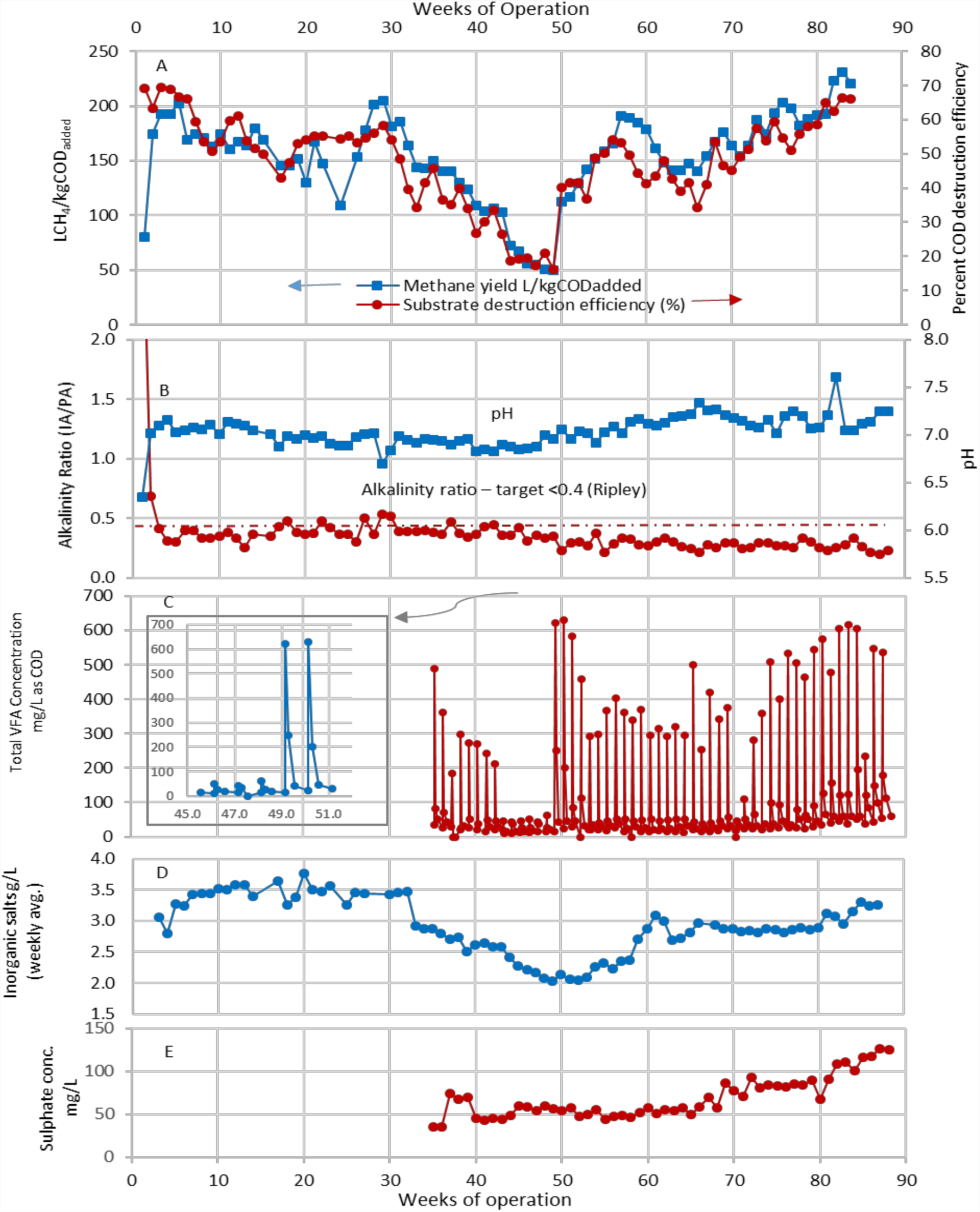
Daisy’s performance vs. time. **A)** weekly methane production (L/kgCOD_added_); substrate COD destr. eff. (%); **B)** alkalinity ratio (wkly avg.) – ideal ratio <0.4, and pH: D) total VFAs, acetate + propionate + butyrate, (mg/L as COD, 4x wkly); **C)** conc. of recirc. inorganic salts (g/L, wkly avg.); **E)** conc. of recirc. sulphate (g/L, wkly avg).

### 3.4 The effect of food waste addition on Daisy’s performance – synergy

One of our main objectives was to study the effects of FW addition on digester performance (Table 1). In Period 3, Daisy was operating at an SRT of 42 days at 17.2%COD_FW_. The average CH_4_ was 278 L.wk^-1^ or 185 L.kg^-1^COD_added_, and substrate destruction efficiency was 54.2%. In Periods 4a, 4b, and 4c, FW addition was reduced in three steps; 12.9%COD_FW_, 7.9%COD_FW_, and 0%COD_FW_, respectively. Each step took 6 weeks (to change all 6 LBs). With each reduction in FW addition CH_4_ production fell, first to 198 L.wk^-1^, then 138 L.wk^-1^, and finally 63 L.wk^-1^ when no food waste was added. Specific CH_4_ production and substrate destruction efficiency also dropped. At each step, CH_4_ production attained its new stable level within 3 weeks. It was immediately apparent that the drop in CH_4_ production could not be accounted for by the reduction in COD_FW_ alone. For example, by Period 4c, FW addition had been reduced by 254 gCOD.wk^-1^, equivalent to 89 L CH_4_.wk^-1^ assuming 100% COD_FW_ conversion, compared to the measured drop of 214 LCH_4._wk ^-1^ This left 125 L CH_4_.wk^-1^ unaccounted for. The apparent explanation was an unreported effect whereby FW addition enhanced the digestibility of the fibres, and that the extent of enhancement was related to the amount of FW added. The objectives of the research were expanded to include investigation of this apparent synergistic effect.

In Period 5, Daisy was returned to 17.2%COD_FW_ in a single step over 42d (six LB changes). After seven weeks (at week 57), CH_4_ production had gradually risen to (a single week value of) 279 L.wk^-1^ and substrate destruction efficiency of 52%. At this point, the supply of BA#4 was running low, so Daisy was switched to BA#5 for 6 weeks. Performance immediately began to decline (Table 1 and Fig 2A); this was provisionally attributed to the physical properties of the particular batch of BA, since no other changes had been made. It took 15 weeks to restore stable CH_4_ output at 245 L.wk^-1^ (169 L.kg^-1^COD _added_) and a COD destruction efficiency of 53.0%. This BA phenomenon is discussed more fully in Section 3.6.

In Period 6a FW addition was raised to 21.2%COD_FW_, CH_4_ production reached 297 L.wk^-1^ or 189 L.kg^-^ 1COD_added_, and a COD destruction efficiency of 56.0%. In Period 6b FW addition was raised once more to 29.3%COD_FW_, CH_4_ production reached 384 L.wk^-1^ or 225 L.kg^-1^COD_added_, and a COD destruction efficiency of 65.3%. In both cases, CH_4_ production increased by an amount greater than could be accounted for by the increase in COD_FW_. The synergistic effect of food waste addition on the digestibility of the lignocellulosic fibres was quantified at each of six levels of COD_FW_ addition, using Equation 11, and plotted in Fig. 3, which also includes substrate destruction efficiency. The magnitude of the synergistic effect is very large and quite obviously related to the amount of FW added. At 29%COD_FW_ the methane yield from the fibre was nearly 3 times the yield at 0%COD_FW_. The data were also plotted as LCH4.kg^-^ 1COD_FBadded_ vs %COD_FW_, (Fig. S4). This shows a very strong linear relationship to the limit of the available data, even when using the more conservative assumption of 100%FW conversion to perform the calculation. It is certain that the effects of FW addition will, at some higher level, become progressively less beneficial, and this needs further study.

**Fig. 3:**
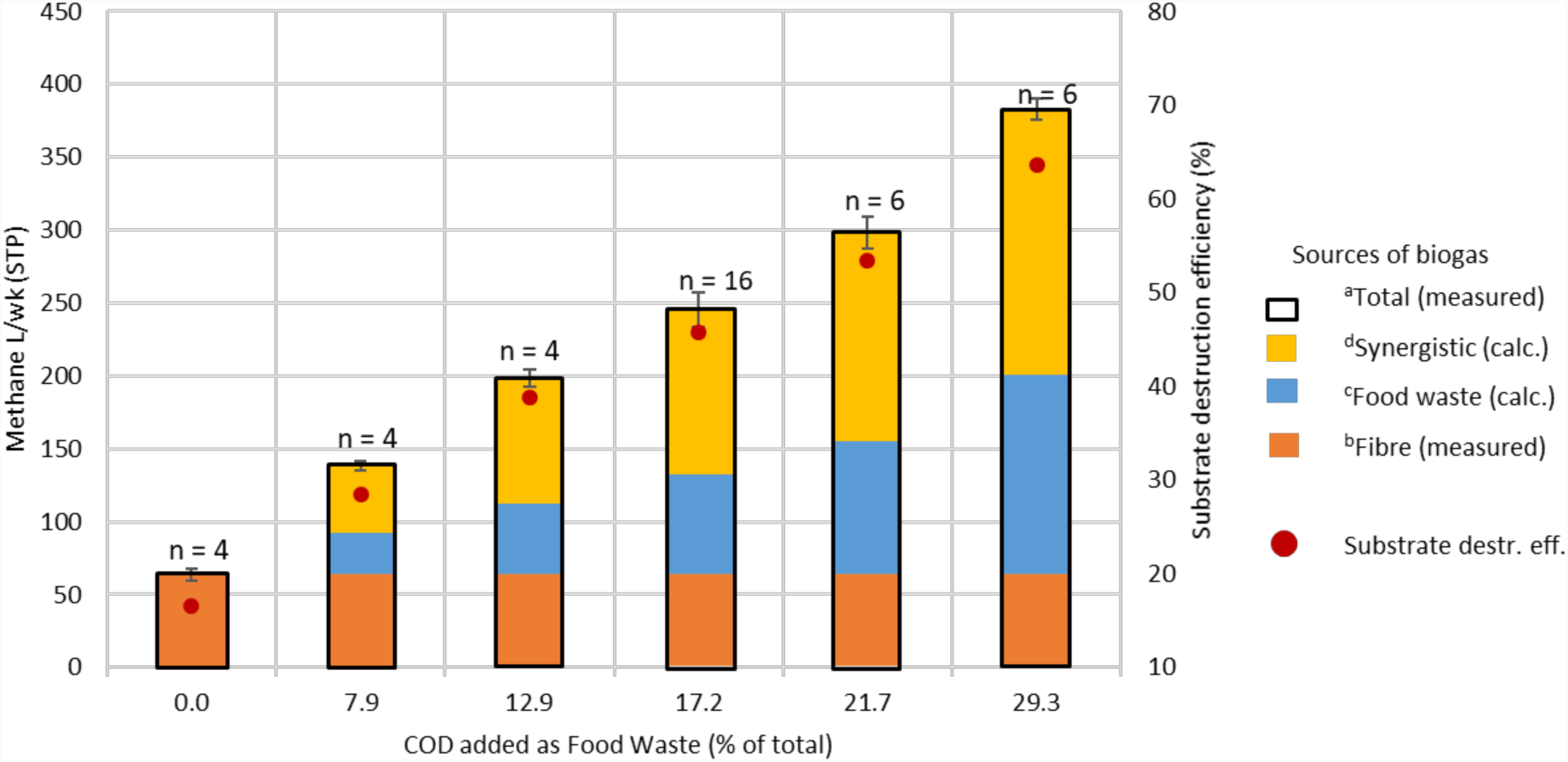
The effect of food waste addition on methane production (bars) and substrate destruction efficiency (red dots). ^a^Total measured vol. CH_4_ (L.wk^-1^); ^b^measured vol. CH_4_ from FB alone (no FW); ^c^calculated vol. CH_4_ from FW added assuming 78% COD conversion; ^d^synergistic biogas from FB as a result of FW addition calculated by difference. All vol. in L.wk^-1^ at STP; methane at 52.4% of biogas and assumed 78% COD_FW_ conv. was determined from BMP tests. (see also Figure S4)

The addition of FW greatly enhanced the digestibility of the FB in direct proportion to the amount of FW added, but to differing degrees for different fibres (see Section 3.5). The mechanism is not entirely clear, but there are indications that it may be enzymatic. Yuan et. al. (2012) subjected samples of FB to microbial pretreatment (for 2 to 10 days) resulting in a doubling of biogas yield. Zhang et. al. (2007) found that extracellular enzymes regulated the hydrolysis of organic waste in a high-solids-content digester. A companion microbiological study conducted on samples of leachate, digestate, and food waste from Daisy show clear trends in microbial abundance related to FW addition (Lee, 2018) and will be reported separately.

### 3.5 The digestibility of individual fibres and bulking agent

Not all the fibres are equally digestible and this offers some further insight into the mechanism of synergy. The digestibility of individual fibre samples embedded in the LBs was assessed using coupon tests. Coupons (tea balls) were present under all operating conditions. The destruction efficiency of all four individual fibres – CB, BB, NP, and FP plus BA, at the same six COD_FW_ addition rates (Table S7), are plotted in Fig. 4. It is immediately apparent that their digestibility is ranked FP>CB>BB>NP>BA and this is consistent with the literature (Buffiere et al., 2008; Eleazer et al., 1997; Pommier et al., 2010). It is also apparent that the differences among them grow wider as %COD_FW_ increases. It would also appear that the digestibility of the fibres may be directly related to the severity of the pulping processes used in their manufacture; FP is chemically pulped and bleached and contains no lignin, CB and BB are also chemically pulped but still contain some lignin (also the latter is coated on one or both sides), NP is mechanically pulped and has a high lignin content, and BA is not pulped at all.

**Fig. 4:**
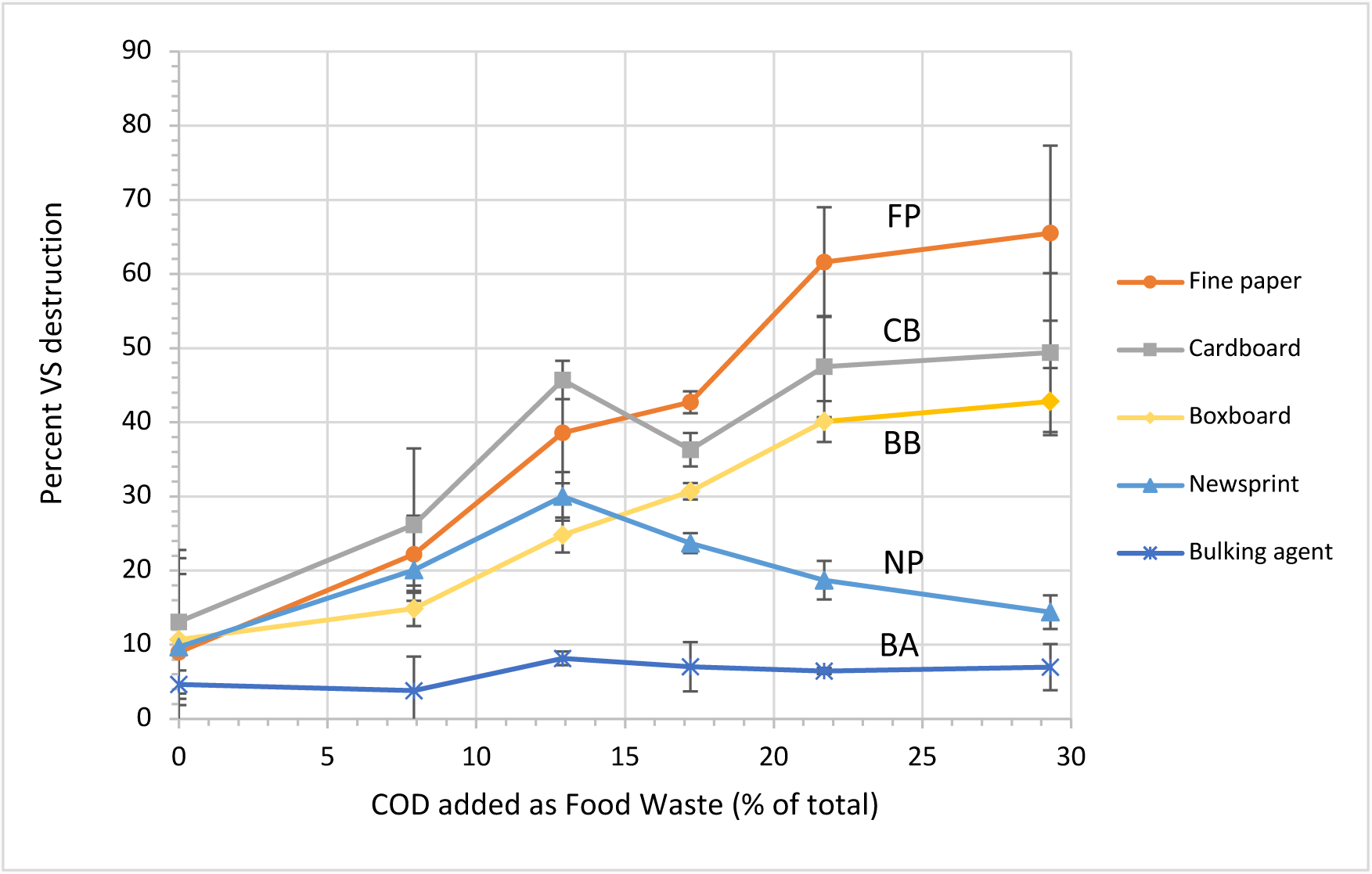
Destruction efficiency of individual FB samples within Daisy vs. FW addition rate based on data from coupon tests. Shows ranking of digestibility FP>CB>BB>NP>BA and effect of %COD_FW_; note absence of FW effect on BA.

The coupon results also provided two further pieces of data; firstly, the digestibility of the BA ranged from 3.8% to 8%, averaged 7.0%, and rose only slightly with FW addition, but the standard deviations are large (Table S7). The average value was used to calculate the amount of undigested COD_BA_, subtract it from the COD_DG_, and calculate substrate destruction efficiency excluding BA. Secondly, it suggested that the digestibility of NP rises, then declines, with FW addition (Fig. 4). This particular anomalous trend for NP requires verification.

### 3.6 The effect of bulking agent on Daisy’s performance

The switch to BA#5 caused a vexing decline in performance (CH_4_ yield and COD destruction efficiency) of about 20%. After six weeks (at week 64), leach beds were progressively switched back to BA#4 (from a reserve supply). Performance gradually improved and, at week 71, the BA was switched again to BA#6, methane production eventually stabilized at prior levels, and Period 6 began at week 74. BA#4 and BA#6 were prepared with a Roto-Chopper (essentially a shredder which produces splinters of wood), and the larger particles screened out; the two batches were similar in appearance and behaviour. BA#5 was very different; it was produced with a chipper, and the particles were coarser, shorter and fatter (Fig. S2). Simple tests of the physical properties of the two batches (Table S8) showed that BA#5 had a slightly higher proportion of coarse particles, about twice the bulk density, and 80% of the water retention capacity of BA#4. The literature shows that digester efficiency is very dependent on maintaining a moisture content of about 80% in SS-AD (Abbassi-Guendouz et al., 2012; Le Hyaric et al., 2012; Motte et al., 2013; Xu et al., 2014). These observations suggest that the physical properties of the BA may play a greater role in digestion efficiency than merely ensuring LB permeability. Another possibility is that the chemical composition of BA#5 was different, perhaps because the wood was greener and recently chipped.

### 3.7 Solids retention time

During Period 1 (weeks 6 to 15) conditions were kept constant in all respects at 17.2%COD_FW_; stable operation was achieved, with an average CH_4_ output of 247 L.wk^-1^ or 169 L.kg^-1^COD_added_, and a substrate destruction efficiency of 53.7% (Table 1 and Fig. 2A). In Period 2 the SRT was extended to 49d from 42d. This had the effect of creating unevenness in L CH_4_.wk^-1^, reflected in the increased coefficient of variation (Table 1). Nevertheless, performance remained unchanged at 172 LCH_4_.kg^-1^COD_added_, and COD destruction efficiency of 53.5%. Extending the SRT to 49d was not beneficial. At week 88, Daisy was shut down and the last six LBs removed simultaneously. Substrate destruction efficiency at 29.3%COD_FW_, was determined for all 6 LBs, and plotted against SRT (Fig. 5). Daisy achieved 68.4% COD destruction at 42d, at which point the curve is almost flat (and presumably close to the asymptote). However, destruction efficiency had already reached 63.5% at 21d and 66.8% at 28d. These results suggest that, at 29.3%COD_FW_, 98% of ultimate performance had already been achieved with an SRT of just 28d. Of necessity, this was a single experiment, but it strongly suggests that the SRT can be significantly reduced with little loss in performance.

**Fig. 5:**
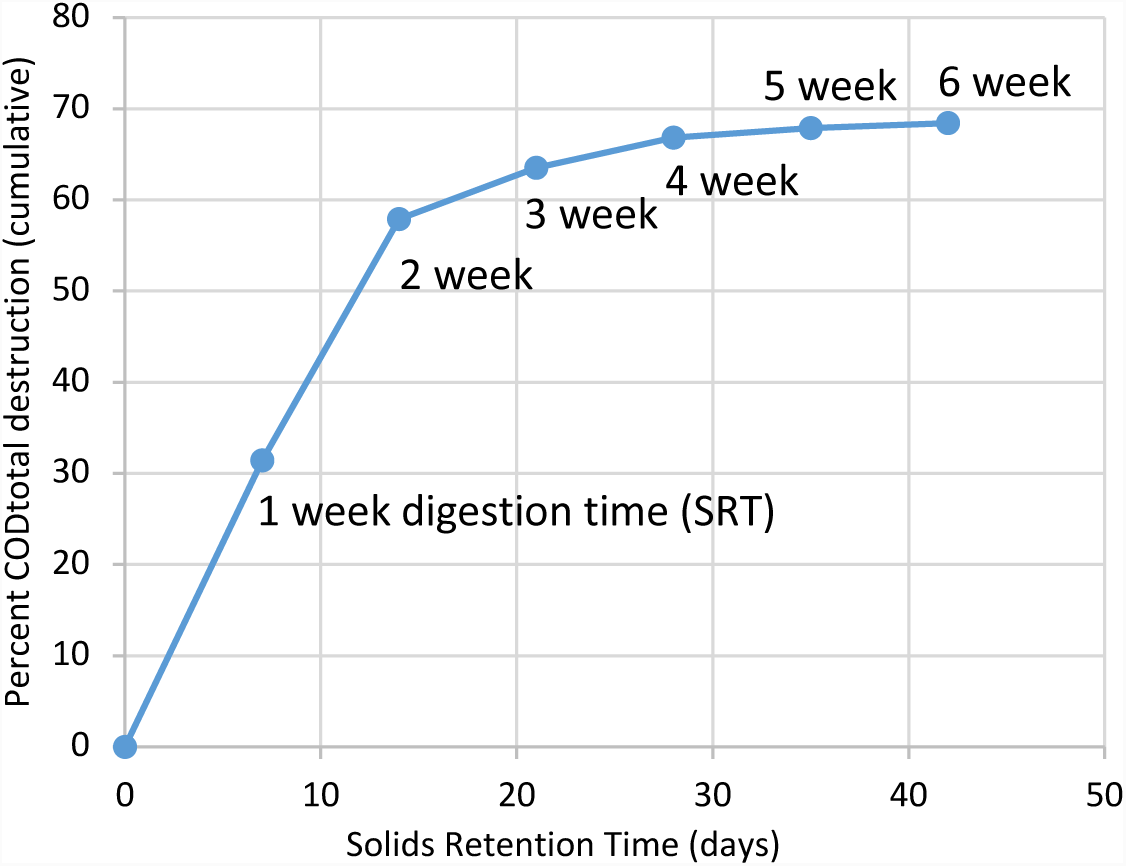
COD Destruction vs Digestion Time at 29%COD_FW._ Data from final week 88 when 6 LBs removed simultaneously (each with a different SRT).

### 3.8 Comparison to other digesters with similar substrates

Daisy’s performance on a VS basis was compared to that of other digesters with similar substrates, (Table 2). Three of the comparators were BMP tests (Eleazer et al., 1997; Pommier et al., 2010; Yuan et al., 2012; Yuan et al., 2014), one was a CSTR (Zhang et al., 2012) and one a comparison of a CSTR to a LB system (Di Maria et al., 2017). All three BMP studies found the same ranking of fibre digestibility as the present research, FP>CB>BB>NP, and all achieved a higher destruction efficiency and biogas yield than Daisy, but all with longer SRTs of 90d, 60d and up to 600d, respectively. At 29.3%COD_FW_ with an SRT of 42d Daisy gave an equivalent performance (296 LCH_4_.kg^-1^VS_added_ and 69%VS_destr._) to a CSTR digesting mechanically-recovered OFMSW (Zhang et al., 2012) with an SRT of 30d (304 LCH_4_.kg^-1^VS_added_ and 62%VS_destr._). Even at an SRT of 28d, Daisy’s performance was virtually undiminished (Fig. 4). Once more, the beneficial effect of FW addition is apparent. Compared to Di Maria et al. (2017), Daisy’s performance surpassed that of their LB system, but was slightly inferior to their CSTR.

**Table 2.** Daisy’s performance compared to that of other digesters fed with similar substrates

| Data Source | Reactor Design and Operation | Substrates(s) | SRT (d) | COD<sub>FW</sub>% | Methane yield mLCH<sub>4</sub>.<br>g<sup>-1</sup>VS<sub>added</sub> | Percent destruction efficiency as VS |
| --- | --- | --- | --- | --- | --- | --- |
| Daisy<br>(this study) | Sequentially-fed leach beds | CB+BB+FP+NP<br>plus FW | 42 | 0 | 66.8 | 20.0 |
|  | plus UASB |  |  | 7.9 | 134.8 | 33.2 |
|  | 6 LBs 50L total |  |  | 12.9 | 184.4 | 43.2 |
|  | UASB 27L |  |  | 17.2 | 218.7 | 56.3 |
|  | 2 Tanks 35L total |  |  | 21.7 | 246.0 | 63.5 |
|  |  |  |  | 29.3 | 296.3 | 69.4 |
| Pommier et al.<br>(2010) | BMP tests, six grams of substrate | CB + BB + NP + FP + magazines | 90 |  | 149.6 | 42.0 |
| Yuan et al.<br>(2012, 2014) | BMP tests: with microbial pre-treatment | CB+FP+NP | 60 |  | 92.9 | N/A |
|  | BMP tests: no microbial pre-treatment |  |  |  | 209.0 | N/A |
| Eleazer et al.<br>(1997) | BMP + daily leachate recirc. | FW | 600 |  | 320.6 | 77.4 |
|  | 2L reactors | NP |  |  | 75.4 | 31.1 |
|  |  | CB |  |  | 155.0 | 54.4 |
|  |  | FP |  |  | 288.0 | 54.6 |
|  |  | MSW |  |  | 122.3 | 58.4 |
| Zhang et al.<br>(2012) | CSTR semi-cont. feed 35L | mr-OFMSW | 30 |  | 304.0 | 62.0 |
| Di Maria et al.<br>(2017) | CSTR - 100L | OFMSW | 30 |  | 320.0 | N/A |
|  | Leach bed - 100L |  |  |  | 252.0 | N/A |

CB = cardboard, BB = boxboard, FP = fine paper, NP = newsprint, FW = food waste, OFMSW = organic fraction of municipal solid waste, mr-OFMSW = mechanically recovered OFMSW. N/A = not available

Overall, this study demonstrated that the operation of a simple solid-state digestion process (Daisy) with no mixing of the solid organic waste remained stable throughout, showing a high tolerance of variations in feedstock, delivered a high methane yield in a much shorter SRT than anticipated, and did so because of the unexpected effect of FW on the digestibility of fibres. This digester design is simple and relatively easily scaled and well suited to the North American situation. Further study is required to determine the limits of synergistic biogas production and its mechanism(s), the effects of SRT and of leachate recirculation rates on digester performance and stability and on the way each LB functions, and the potential to operate without the UASB. The rising concentration of sulphate, in response to increased food waste addition, raises questions about the apparent lack or suppression of sulphate reducing bacteria which also requires investigation. The unexpectedly strong performance of Daisy suggests larger scale demonstrations should be undertaken.

## ACKNOWLEDGMENTS

This research was funded by the Natural Sciences and Engineering Research Council of Canada (Collaborative Research and Development Grant), and by Miller Waste Systems Inc. We thank Mike Kopansky, John Tomory and Charlie Cassin of Miller Waste Systems for preparing and providing samples, our colleagues at the University of Toronto, Brad Saville, Grant Allen, Savia Gavazza, Paul Jowlabar, Glenn Wilson, Greg Brown and Andy Quaile, for their experience and insights.

## SUPPORTING INFORMATION

Detailed description of analytical methods, eight supplementary tables of raw data and calculations, and four supplementary figures showing coupon placement, bulking agent batches and specific methane production.

## REFERENCES (formatted as author-date for review purposes only)

Abbassi-Guendouz, A., Brockmann, D., Trably, E., Dumas, C., Delgenes, J.P., Steyer, J.P., Escudie, R. 2012. Total solids content drives high solid anaerobic digestion via mass transfer limitation. Bioresource Technology, 111, 55–61.

APHA. 1992. Standard methods for the examination of water and wastewater. 18th ed, American Public Health Association. Washington DC.

Browne, J.D., Allen, E., Murphy, J.D. 2013a. Improving hydrolysis of food waste in a leach bed reactor. Waste Management, 33(11), 2470–2477.

Browne, J.D., Murphy, J.D. 2013b. Assessment of the resource associated with biomethane from food waste. Applied Energy, 104, 170–177.

Browne, J.D., Murphy, J.D. 2014. The impact of increasing organic loading in two phase digestion of food waste. Renewable Energy, 71, 69–76.

Buffiere, P., Frederic, S., Marty, B., Delgenes, J.P. 2008. A comprehensive method for organic matter characterization in solid wastes in view of assessing their anaerobic biodegradability. Water Science and Technology, 58(9), 1783–1788.

City of Ottawa. 2007. ICI and CD Management Options Report: ICI 3Rs Strategy Project. Ottawa.

De Baere, L., Mattheeuws, B. 2013. Anaerobic digestion of the organic fraction of municipal solid waste in Europe – Status, experience and prospects. in: Waste Management, (Eds.) K.J. Thomé-Kozmiensky, S. Thiel, pp. 517–526.

Di Maria, F., Barratta, M., Bianconi, F., Placidi, P., Passeri, D. 2017. Solid anaerobic digestion batch with liquid digestate recirculation and wet anaerobic digestion of organic waste: Comparison of system performances and identification of microbial guilds. Waste Management, 59, 172–180.

Eleazer, W.E., Odle, W.S., Wang, Y.S., Barlaz, M.A. 1997. Biodegradability of municipal solid waste components in laboratory-scale landfills. Environmental Science & Technology, 31(3), 911–917.

Environment Canada. 2017. National Inventory Report 1990-2015 Greenhouse Gas Sources and Sinks in Canada, Queen’s Printer. Ottawa.

European Union. 1999. European Union Landfill Directive 1999/31/EC, April 26, 1999. European Union Publications Office.

Forrestal, B.J., Guilford, N.G.H., Poland, R.J. 2006a. Canadian Patent 2,468,158; System and Method for the Production of Biogas and Compost, BioPower Energy Inc. Canada.

Forrestal, B.J., Guilford, N.G.H., Poland, R.J. 2006b. United States Patent 7,101,481; System for the Production of Biogas and Compost from Organic Materials and Method of Operating an Organic Treatment Facility, BioPower Energy Inc. USA.

Government of Canada. 2010. Environment Canada – Report on Waste Management, Environment Canada. Ottawa.

Government of Canada. 2015. State of Waste Management in Canada, Queen’s Printer. Ottawa.

Government of Ontario. 2004. Ontario’s 60% Waste Diversion Goal – A Discussion Paper, (Ed.) Ministry of the Environment, Queen’s Printer for Ontario. Toronto.

Guilford, N.G.H. 2017. The Anaerobic Digestion of Organic Solid Wastes of Variable Composition. PhD Thesis, Department of Chemical Engineering and Applied Chemistry, University of Toronto. Toronto, pp. 240. https://tspace.library.utoronto.ca/handle/1807/80954

Guilford, N.G.H. 2009. A New technology for the Anaerobic Digestion of Organic Waste. M.Eng thesis, Department of Chemical Engineering and Applied Chemistry, University of Toronto. Toronto, pp. 112. https://tspace.library.utoronto.ca/handle/1807/18314

Hodge, K.L., Levis, J.W., DeCarolis, J.F., Barlaz, M.A. 2016. Systematic Evaluation of Industrial, Commercial, and Institutional Food Waste Management Strategies in the United States. Environmental Science & Technology, 50(16), 8444–8452.

Igoni, A.H., Ayotamuno, M.J., Eze, C.L., Ogaji, S.O.T., Probert, S.D. 2008. Designs of anaerobic digesters for producing biogas from municipal solid-waste. Applied Energy, 85(6), 430–438.

Le Hyaric, R., Benbelkacem, H., Bollon, J., Bayard, R., Escudie, R., Buffiere, P. 2012. Influence of moisture content on the specific methanogenic activity of dry mesophilic municipal solid waste digestate. Journal of Chemical Technology and Biotechnology, 87(7), 1032–1035.

Lee, HW. 2018. Microbial Diversity and Abundance in a Sequentially-fed Anaerobic Digester treating Food Waste and Lignocellulosic fibres. MASc thesis, Department of Chemical Engineering and Applied Chemistry, University of Toronto. https://tspace.library.utoronto.ca/handle/1807/89607

Motte, J.C., Escudie, R., Bernet, N., Delgenes, J.P., Steyer, J.P., Dumas, C. 2013. Dynamic effect of total solid content, low substrate/inoculum ratio and particle size on solid-state anaerobic digestion. Bioresource Technology, 144, 141–148.

Murto, M., Bjornsson, L., Rosqvist, H., Bohn, I. 2013. Evaluating the biogas potential of the dry fraction from pretreatment of food waste from households. Waste Management, 33(5), 1282–1289.

Pommier, S., Llamas, A.M., Lefebvre, X. 2010. Analysis of the outcome of shredding pretreatment on the anaerobic biodegradability of paper and cardboard materials. Bioresource Technology, 101(2), 463–468.

Rittmann, B.E., McCarty, P.L. 2001. Environmental Biotechnology: Principles and Applications. McGraw-Hill.

Satchwel, A.J., Scown, C.D., Smith, S.J., Arnirebrahimi, J., Jin, L., Kirchstetter, T.W., Brown, N.J., Preble, C.V. 2018. Accelerating the Deployment of Anaerobic Digestion to Meet Zero Waste Goals. Environmental Science & Technology, 52(23), 13663–13669.

Wu, X., Yao, W.Y., Zhu, J., Miller, C. 2010. Biogas and CH4 productivity by co-digesting swine manure with three crop residues as an external carbon source. Bioresource Technology, 101(11), 4042–4047.

Xu, F.Q., Wang, Z.W., Tang, L., Li, Y.B. 2014. A mass diffusion-based interpretation of the effect of total solids content on solid-state anaerobic digestion of cellulosic biomass. Bioresource Technology, 167, 178–185.

Yadvika, S., T.R. Sreekrishnan, Sangeeta Kohli, Vineet Rana. 2004. Enhancement of biogas production from solid substrates using different techniques. Bioresource Technology, 95, 1–10.

Yuan, X.F., Cao, Y.Z., Li, J.J., Wen, B.T., Zhu, W.B., Wang, X.F., Cui, Z.J. 2012. Effect of pretreatment by a microbial consortium on methane production of waste paper and cardboard. Bioresource Technology, 118, 281–288.

Yuan, X.F., Wen, B.T., Ma, X.G., Zhu, W.B., Wang, X.F., Chen, S.J., Cui, Z.J. 2014. Enhancing the anaerobic digestion of lignocellulose of municipal solid waste using a microbial pretreatment method. Bioresource Technology, 154, 1–9.

Zhang, B., He, P.J., Lu, F., Shao, L.M., Wang, P. 2007. Extracellular enzyme activities during regulated hydrolysis of high-solid organic wastes. Water Research, 41(19), 4468–4478.

Zhang, Y., Banks, C.J., Heaven, S. 2012. Anaerobic digestion of two biodegradable municipal waste streams. Journal of Environmental Management, 104, 166–174.

