## Supplemental Information File for "Solid state anaerobic digestion of mixed organic waste: the synergistic effect of food waste addition on the destruction of paper and cardboard"

Number of Pages: 16

Number of Tables: 8

Number of Figures: 4

**TABLE OF CONTENTS**

**Text**

Details of sampling and analytical methods: pages 3-4

Description of Supplementary Tables and Figures: pages 4-5

**List of Supplementary Tables**

**Table S1**. Table of data recorded for each leach bed by serial number

**Table S2.** Substrate and digestate analysis

**Table S3.** Digestate properties: wet wt., TS, VS and COD for all 87 LBs

**Table S4.** Biogas methane content

**Table S5.** Biogas and methane production by week

**Table S6.** Mass balance by period

**Table S7.** Substrate destruction efficiency (as VS), by substrate, vs $COD_{FW}$ addition – from coupon data

**Table S8.** Physical properties of bulking agents BA#4 and BA#5

**List of Supplementary Figures**

**Figure S1**. Triplicate coupons being embedded in the waste mass

**Figure S2**. Bulking agent a) BA#4; b) BA#5; BA#5 coarser and of higher bulk density

**Figure S3.** Stoichiometry of digestion of the 83 week weighted average substrate and the resultant percent methane in biogas, calculated from first principles

**Figure S4**. $CH_4$ produced/kg $COD_{FBadded}$ vs. percent $COD_{FWadded}$ (using Equation 11); result at 100% $COD_{FW}$ conversion included for comparison

**Details of Sampling and Analytical Methods**

The elemental composition of each individual component of the feedstock was determined using a Thermoflash 2000 CHN analyzer; also by TKN to achieve greater precision for nitrogen. Total solids (TS), volatile solids (VS) and chemical oxygen demand (COD) were measured using standard methods (APHA, 1992); VS is simple to measure, and more useful for comparison with published results, most of which express performance in terms of VS destruction; the COD measurements formed the basis of the mass balance calculation. Sample sizes ranged from 1 to 6g on a dry basis. A dry method for sample preparation, using a Wiley mill, was devised to create homogeneous samples for COD measurement of FW, FB, and BA. Each sample was dried and processed through a Wiley mill with a 60 mesh (0.25mm) screen (or a 40 mesh (0.42mm) screen for the FW and NP, because of their tendency to plug the finer screen). The milled samples were then suspended in water, diluted and pipetted, to achieve a COD concentration within the range of the spectrophotometer. Samples were analyzed in sextuplicate. The COD content of all feedstocks was also calculated from their elemental composition, for comparison to the measured values (Table S2). In all cases, the initial sample size was 25g for FB, BA and DG and 250g for FW (all on an as-received basis) and 20g for digestate on a wet basis.

The physical properties of the bulking agent batches, BA#4 and BA#5, were compared in an attempt to explain changes in Daisy's performance in Period 5. Duplicate dry samples BA were screened at >3.36mm > 0.5mm > 0.21mm and <0.21mm. Bulk density was measured by weighing 1L beakers of loose, dry, BA. Water absorption capacity was measured by weighing 50g of BA into a 1L beaker, adding 400 mL of deionized water, covering the beakers and leaving them for 3 days at room temperature. The contents were then discharged onto a 0.21mm screen and left to drain for 1h. The amount of retained water, in $gH_2O.gTS^{-1}$ was then calculated.

Two 50 mL samples of leachate were withdrawn from valve V2 (Fig. 1) four times per week: immediately prior to the leach bed exchange, 6h later, 1d later and 3d later (an equal volume of deionized (DI) water was returned to Daisy). One 50 mL sample was analyzed for TS, VS, and COD using standard methods (APHA 92), and the second 50 mL sample to measure the pH and alkalinity ratio, using a pH meter and a two-step titration (Ripley et al., 1986) respectively. The residue from VS determinations was also used to calculate the concentration of inorganic salts recirculating within Daisy.

From week 35 onwards two additional 10 mL samples were taken four times per week. The first was prepared and stored for subsequent microbial analysis; samples were centrifuged at 7000g; approximately 9 mL of supernatant was collected and stored at 4°C for soluble COD analysis; the pellet was re-suspended in the remaining supernatant and centrifuged at 10,000g for 15 minutes; the supernatant was discarded and the pellet stored at -80°C for subsequent DNA extraction and analysis. The second 10 mL sample was filtered using 0.22µm nylon syringe filter and stored at -20°C for subsequent ion chromatography (IC) analysis.

VFAs and sulphate were analyzed using a Dionex ICS-2100 (IC) with an RFIC (reagent-free ion chromatography) Ionpac with AG18 guard column/AS18 analytical column. Runtime was 23 minutes per sample. The eluent was KOH solution at 1mL/min. Using the gradient method, the eluent concentration was increased from 2mM to 32mM over 15 min., held at 32mM for 3 min., then reduced to 2mM for 5 min. Suppressor values were 80mA. A six-point calibration was used in every run; two blanks were run prior to, and following, each set of standards.

Samples of biogas (200 µL) were extracted through septa on the infeed lines to GM1 and GM2 using a gastight syringe, approximately every two weeks, and analyzed for $CH_4$ and $CO_2$ using a Hewlett Packard 5890 gas chromatograph, equipped with a CTR I column and a thermal conductivity detector (TCD). Helium was used as the carrier gas at a column pressure of 180 kPa. The injector and detector were both set at 200°C. The oven was operated isothermally at 50°C and the run time was 11 min. The GC was calibrated 3 times during the course of the experiment. The $CH_4$ and $CO_2$ content were determined from the calibration curves. The results were averaged for each GM, and the standard deviation calculated.

Temperatures (every 15 minutes) and biogas volumes (every 5 min. and every hour) were recorded in the data logger, downloaded daily, converted to Excel format, and plotted against time to give an immediate picture of the conditions within Daisy. The datalogger was also checked in real time, early morning and late at night; occasionally necessitating a trip to the lab to take corrective action. A daily activity log was maintained, with a link to the data logger output, to record all inputs and outputs (time, type and volume), system adjustments, operating anomalies, and corrective measures.

**Description of Supplementary Tables and Figures**

**Table S1** is an example of the data recorded for each of the 87 leach beds installed, by Serial #. It describes the sources and dates of delivery of the waste samples, how they were prepared, the quantities used, the amount of water added. It records the gross and tare weights of the leach beds before and after digestion, corrections for the weight of the coupons, and the measured headspace before and after. Leach bed purging and pressurization data are also recorded.

**Table S2** summarizes the average properties of each of the substrates; these include elemental composition, ash content, the calculated and measured COD content, and the measured VS content. It also includes the average COD and VS content of the digestate. These data were used to calculate the CHON stoichiometric formula of each substrate, and the overall stoichiometric equation of AD (Fig. S3) for the 83-week average substrate and the percent $CH_4$ in the biogas generated. Note the close correspondence between the measured and calculated COD values. The mass balance was based on measured COD averages, individual $COD_{DG}$ measurements, and average $CH_4$ content.

**Table S3** contains the wet weight, total solids, volatile solids and COD content of the digestate recovered from all 87 leach beds. The $COD_{DG}$ for the first 11 leach beds was not measured; the values presented are calculated from the measured VS and average VS/COD ratio for the other 76 leach beds.

**Table S4** summarizes the biogas analytical data, showing the higher $CH_4$ content of the biogas produced in the UASB (GM1) at 58.5%, compared to that from the balance of the system (GM2) at 51.7%. It also shows that the majority of the $CH_4$ (88%) was produced in the leach beds and tanks (GM2). The overall average $CH_4$ content was measured at 52.4%.

**Table S5** contains the weekly biogas and $CH_4$ production data, as measured and corrected to STP.

**Table S6** summarizes the mass balance by operating period using Methods A and B (Equations 6A and 6B), as described in Section 3.2; Method A shows slightly greater variability between periods than Method B. Under changing conditions (as distinct from stable operations with constant feed composition) it is almost impossible to correlate weekly $CH_4$ production which comes from the entirety of the system (6 LBs, two tanks, and the UASB) to substrate destruction, which is measured in the single LB removed in that same week. In reality, that single LB made its contribution to gas generation over 6 weeks, with most of it being generated in the first four weeks (Fig. 4), while its substrate destruction efficiency is recorded in a single week. Equation A is mathematically more sensitive to this effect than Equation B.

**Table S7** presents the effect of FW addition on the digestibility, on a VS basis, of individual substrates (CB, BB, NP, FP and BA), as measured by the coupon method. These results are also presented in Fig. 5.

**Table S8** compares the physical properties of bulking agent batches BA#4 and BA#5 measured as particle size distribution, bulk density and water absorption capacity.

**Figure S1** shows the placement of one triplicate set of coupon samples of a single fibre in the middle of a leach bed. This leach bed will also contain a second triplicate set, placed in the waste above the one shown.

**Figure S2** shows the difference in morphology of the two bulking agent samples BA#4 and BA#5, the latter being noticeably shorter and fatter.

**Figure S3** is the calculation, from first principles, of the stoichiometry of the digestion of the average waste fed to Daisy over 83 weeks, and the calculation of the $CH_4$ content of the biogas at 52.5% (to be comparex to the measured value of 52.4% in Table S4).

**Figure S4** provides $CH_4$ produced/kg $COD_{FBadded}$ vs. percent $COD_{FWadded}$ (using Equation 11) assuming 100%$COD_{FW}$ conversion. This figure is included for comparison with Figure 3 in the main text at 78%$COD_{FW}$ conversion

**Table S1: Example of data recorded for each Leach bed; Example of Serial Number S.071**

| | | | | | | |
|---|---|---|---|---|---|---|
| Serial Number: | S.071 | | | | | |
| **A. General** | | | | | | |
| LB number: | LB01 | | | | | |
| LL number: | LL05 | | | | | |
| Date and Time in: | Aug-3-16 | 10.21h | | | | |
| Date and Time out: | Sep-13-16 | 11.35h | | | | |
| **B. System set-up** | | | | | | |
| Geotextile bottom | Mirafi FW 400 | | | | | |
| Geotextile top | Mirafi 1600N | | | | | |
| Overflow tube (Y/N) | Y | | | | | |
| RTD number. | RTD06 | | | | | |
| **C. Contents** | Code | Source | Date received | Weight (g) | Comments | |
| (i) Fibre | CB | Miller | Mar 2 2016 | 440 | 4th shipment | |
| | BB | Miller | Mar 2 2016 | 350 | 4th shipment | |
| | NP | Miller | Mar 2 2016 | 110 | 2nd shipment | |
| | FP | Wallberg | May 10 2014 | 200 | | |
| (ii) Foodwaste | FW | Miller | May 1 2016 | 675 | 23rd LB at 38%FW | |
| | | | | | 3rd LB with FW2 | |
| | | | | | blended | |
| (iii) Leaf and yard waste | LY | | | nil | | |
| (iv) Bulking agent | BA | Miller | Jul-16 | 500 | dry | 3rd LB with BA6 |
| (v) Water | W | | | 3800 | | |
| (vi) Inoculum | IN | | | nil | | |
| Total | | | | 6075 | | |

**Table S1: Example of data recorded for each Leach bed; Example of Serial Number S.071 (Cont'd)**

| | | | | | | |
|---|---|---|---|---|---|---|
| **D. Leach Bed Weight (kg)** | Day | Gross | Net | | | Comments |
| Tare: | Zero | 12.385 | 6.005 | 5.916 | 70 | Difference |
| 6.38 | 43 | 11.520 | 5.140 | 5.075 | 65 | Water in Geotextile |
| | | | -6.380 | | | |
| **E. Samples** | Date | Code | Quantity (g or ml)) | Storage | Comments | |
| **F. Coupons** | Container | | Sample | | | |
| | Type | Code | Gross weight in (g) | Gross wt out (g) | Location | Comments |
| | SSsmL | FP/BB | 88.7 | 124.3 | -20/-25cm | |
| **G. Freeboard** | Day | Height (cm) | | | | |
| | Zero | 8.3 | | | | |
| | 43 | 9.5 | | | | |
| **H. Purge and Pressurize** | Gas | Purge time (mins) | ΔP (cm $H_2O$) | Gas meter pressure | | Comments |
| | Ar | 5 | 61 | GM1 = 12.5 cm | GM2 = 11.5 cm | topped up GM1 |
| **I. Removal Sequence** | | | | | | |
| | Time | Vol. Leachate | Weight kg | Comments | | |
| (ii) Drainage (initial) | | | | | | |
| (iii) Drainage (intermed.) | | | 11.77 | | | |
| (i) Drainage (total) | 24h | 250 | 11.52 | | | |
| **J. General Observations** | | | | | | |
| T1 = +10mm | T2 = +5mm | | | | | |

### Table S2: Substrate and Digestate Analysis

| Sample | Elemental Analysis (wt. %)[a] | | | | | | Calculated COD[b] | | Measured COD[c] | | Measured VS[c] | |
|---|---|---|---|---|---|---|---|---|---|---|---|---|
| | C | H | O | N | Ash | Stoichiometric formula | gCOD /gTS | n | gCOD /gTS | n | gVS /gTS | n |
| Cardboard (CB) | 41.3 ±1.0 | 5.96 ±0.16 | 44.1 ±0.8 | 0.17 ±0.01 | 8.55 ±0.67 | $C_{284}H_{508}O_{235}N$ | 1.13 ±0.03 | 3 | 1.14 ±0.05 | 4 | 0.91 ±0.01 | 6 |
| Boxboard (BB) | 39.4 ±1.1 | 5.67 ±0.11 | 42.2 ±0.4 | 0.11 ±0.003 | 12.7 ±1.4 | $C_{407}H_{730}O_{332}N$ | 1.10 ±0.02 | 3 | 1.13 ±0.02 | 4 | 0.87 ±0.01 | 6 |
| Newsprint (NP) | 46.8 ±1.3 | 6.39 ±0.23 | 45.8 ±2.0 | 0.06 ±0.003 | 0.94 ±0.95 | $C_{865}H_{1474}O_{668}N$ | 1.30 ±0.04 | 3 | 1.33 ±0.02 | 4 | 0.99 ±0.00 | 6 |
| Fine paper (FP) | 37.2 ±0.9 | 5.57 ±0.16 | 41.0 ±2.2 | 0.03 ±0.01 | 16.3 ±2.27 | $C_{989}H_{1781}O_{808}N$ | 1.05 ±0.03 | 3 | 1.00 ±0.03 | 4 | 0.85 ±0.02 | 6 |
| Food waste (FW) | 47.7 ±2.0 | 6.92 ±0.08 | 32.4 ±2.4 | 3.04 ±0.34 | 9.97 ±0.14 | $C_{17.0}H_{29.0}O_{8.3}N$ | 1.46 ±0.07 | 3 | 1.30 ±0.11 | 1 6 | 0.89 ±0.01 | 11 |
| Bulking agent (BA) | 46.6 ±0.6 | 6.19 ±0.07 | 45.1 ±0.8 | 0.25 ±0.12 | 1.84 ±0.67 | $C_{370}H_{591}O_{262}N$ | 1.28 ±0.02 | 3 | 1.29 ±0.05 | 5 | 0.98 ±0.01 | 16 |
| Digestate (DG) | | | | | | | | | [c]1.17 ±0.04 | 8 7 | 0.86 ±0.03 | 87 |

[a]n = no. of samples for elemental analysis and calculated COD. Results all ±1SD. C, H, measured using thermoflash 2000CHN analyzer; N measured using TKN method; ash measured as residue of VS analysis; O calculated by difference. [b]COD calculated from first principles using elemental composition; [b]COD and VS measured using standard methods. [c] Measured COD averages used for mass balance inputs, and individual $COD_{DG}$ measurements and average $CH_4$ content used for mass balance outputs. Individual $COD_{DG}$ measurements are found in Table S3.

### Table S3: Digestate (DG) properties: wet wt., TS, VS and COD for all 87 LBs

| LB Serial # | Period | Wet wt. DG g | Dry wt. DG g | g TS/g DG | g VS/g DG | [a]g COD/g DG |
|---|---|---|---|---|---|---|
| 1 | 1 | 4830 | 1024 | 0.21 | 0.89 | *1.21* |
| 2 | | 4190 | 888 | 0.21 | 0.86 | *1.18* |
| 3 | | 4395 | 945 | 0.22 | 0.87 | *1.19* |
| 4 | | 4525 | 941 | 0.21 | 0.81 | *1.10* |
| 5 | | 4530 | 929 | 0.21 | 0.82 | *1.12* |
| 6 | | 4500 | 950 | 0.21 | 0.83 | *1.13* |
| 7 | | 4720 | 963 | 0.20 | 0.83 | *1.13* |
| 8 | | 4685 | 1021 | 0.22 | 0.85 | *1.16* |
| 9 | | 5145 | 1101 | 0.21 | 0.86 | *1.16* |
| 10 | | 5180 | 1134 | 0.22 | 0.86 | *1.16* |
| 11 | | 5058 | 1087 | 0.22 | 0.86 | *1.17* |
| 12 | | 4942 | 1043 | 0.21 | 0.83 | 1.13 |
| 13 | | 4855 | 1015 | 0.21 | 0.84 | 1.15 |
| 14 | | 5135 | 1109 | 0.22 | 0.84 | 1.14 |
| 15 | | 5095 | 1121 | 0.22 | 0.85 | 1.16 |
| 16 | | 5092 | 1141 | 0.22 | 0.85 | 1.16 |

**Table S3: Digestate (DG) properties: wet wt., TS, VS and COD for all 87 LBs**

| | | | | | | |
|---|---|---|---|---|---|---|
| 17 | 2 | 5294 | 1223 | 0.23 | 0.86 | 1.17 |
| 18 | | 5224 | 1170 | 0.22 | 0.86 | 1.17 |
| 19 | | 4947 | 1088 | 0.22 | 0.86 | 1.18 |
| 20 | | 4877 | 1107 | 0.23 | 0.85 | 1.14 |
| 21 | | 4924 | 1113 | 0.23 | 0.81 | 1.12 |
| 22 | | 4808 | 1106 | 0.23 | 0.83 | 1.13 |
| 23 | | 4760 | 1081 | 0.23 | 0.82 | 1.17 |
| 24 | | 4685 | 1068 | 0.23 | 0.82 | 1.17 |
| 25 | 3 | 4625 | 1101 | 0.24 | 0.81 | 1.16 |
| 26 | | 4798 | 1084 | 0.23 | 0.82 | 1.16 |
| 27 | | 4527 | 1064 | 0.23 | 0.83 | 1.16 |
| 28 | | 4639 | 1062 | 0.23 | 0.83 | 1.13 |
| 29 | | 4950 | 1119 | 0.23 | 0.84 | 1.13 |
| 30 | | 5021 | 1180 | 0.24 | 0.84 | 1.14 |
| 31 | 4a | 5213 | 1220 | 0.23 | 0.87 | 1.17 |
| 32 | | 5195 | 1226 | 0.24 | 0.86 | 1.22 |
| 33 | | 5115 | 1192 | 0.23 | 0.86 | 1.17 |
| 34 | | 5030 | 1202 | 0.24 | 0.86 | 1.13 |
| 35 | | 5157 | 1253 | 0.24 | 0.87 | 1.17 |
| 36 | | 5239 | 1278 | 0.24 | 0.86 | 1.16 |
| 37 | 4b | 5103 | 1240 | 0.24 | 0.87 | 1.13 |
| 38 | | 5055 | 1254 | 0.25 | 0.87 | 1.18 |
| 39 | | 5098 | 1346 | 0.26 | 0.90 | 1.17 |
| 40 | | 5078 | 1300 | 0.26 | 0.89 | 1.18 |
| 41 | | 5124 | 1281 | 0.25 | 0.89 | 1.16 |
| 42 | | 5089 | 1349 | 0.27 | 0.91 | 1.17 |
| 43 | 4c | 4925 | 1359 | 0.28 | 0.91 | 1.17 |
| 44 | | 4910 | 1365 | 0.28 | 0.90 | 1.16 |
| 45 | | 4952 | 1367 | 0.28 | 0.89 | 1.16 |
| 46 | | 4901 | 1372 | 0.28 | 0.91 | 1.17 |
| 47 | | 4900 | 1313 | 0.27 | 0.91 | 1.19 |
| 48 | | 4920 | 1343 | 0.27 | 0.91 | 1.21 |
| 49 | 5a | 5239 | 1226 | 0.23 | 0.89 | 1.20 |
| 50 | | 5313 | 1206 | 0.23 | 0.91 | 1.20 |
| 51 | | 5133 | 1186 | 0.23 | 0.90 | 1.22 |
| 52 | | 5165 | 1224 | 0.24 | 0.90 | 1.24 |
| 53 | | 5059 | 1159 | 0.23 | 0.90 | 1.16 |
| 54 | | 4999 | 1130 | 0.23 | 0.87 | 1.17 |
| 55 | | 5010 | 1062 | 0.21 | 0.86 | 1.19 |
| 56 | | 4970 | 1118 | 0.23 | 0.88 | 1.14 |
| 57 | 5b | 4703 | 1133 | 0.24 | 0.88 | 1.17 |
| 58 | | 4684 | 1134 | 0.24 | 0.89 | 1.24 |
| 59 | | 4880 | 1200 | 0.25 | 0.89 | 1.21 |
| 60 | | 4847 | 1183 | 0.24 | 0.89 | 1.20 |
| 61 | | 4977 | 1199 | 0.24 | 0.89 | 1.13 |
| 62 | | 4865 | 1192 | 0.25 | 0.89 | 1.20 |
| 63 | 5c | 5163 | 1229 | 0.24 | 0.88 | 1.21 |
| 64 | | 5232 | 1188 | 0.23 | 0.87 | 1.22 |
| 65 | | 5203 | 1259 | 0.24 | 0.87 | 1.23 |
| 66 | | 5409 | 1233 | 0.23 | 0.88 | 1.18 |
| 67 | | 4981 | 1121 | 0.22 | 0.87 | 1.14 |
| 68 | | 5046 | 1110 | 0.22 | 0.87 | 1.24 |
| 69 | 5d | 5095 | 1202 | 0.24 | 0.86 | 1.16 |
| 70 | | 5021 | 1155 | 0.23 | 0.85 | 1.16 |
| 71 | | 5075 | 1142 | 0.23 | 0.86 | 1.14 |
| 72 | | 4725 | 1073 | 0.23 | 0.85 | 1.13 |
| 73 | | 4879 | 1069 | 0.22 | 0.85 | 1.19 |
| 74 | | 4664 | 1035 | 0.22 | 0.85 | 1.18 |

**Table S3: Digestate (DG) properties: wet wt., TS, VS and COD for all 87 LBs**

| | | | | | | |
|---|---|---|---|---|---|---|
| 75 | 6a | 4743 | 1058 | 0.22 | 0.84 | 1.23 |
| 76 | | 4902 | 1108 | 0.23 | 0.83 | 1.22 |
| 77 | | 4925 | 1079 | 0.22 | 0.84 | 1.19 |
| 78 | | 4730 | 1055 | 0.22 | 0.81 | 1.18 |
| 79 | | 4613 | 1047 | 0.23 | 0.84 | 1.18 |
| 80 | 6b | 4742 | 977 | 0.21 | 0.84 | 1.20 |
| 81 | 6b | 4765 | 991 | 0.21 | 0.84 | 1.23 |
| 82 | | 4631 | 1028 | 0.22 | 0.84 | 1.12 |
| 83 | | 4726 | 1026 | 0.22 | 0.83 | 1.13 |
| 84 | | 4799 | 1061 | 0.22 | 0.85 | 1.11 |
| 85 | | 4834 | 1112 | 0.23 | 0.83 | 1.11 |
| 86 | | 4365 | 1065 | 0.24 | 0.86 | 1.25 |
| 87 | | 5496 | 1401 | 0.26 | 0.85 | 1.27 |

[a]COD values in *italics* (Serial #s 1 to 11) were calculated from average VS/COD ratio of Serial #s 12 to 87

**Table S4. Biogas methane content**

| Source | %$CH_4$ | STDEV | n | [a]Percentage of total $CH_4$ |
|---|---|---|---|---|
| GM1 | 58.5 | 3.7 | 37 | 12.2 |
| GM2 | 51.7 | 3.6 | 40 | 87.8 |
| Weighted avg. = 52.4 % CH4 | | | | |

[a]fraction of methane produced in UASB (GM1) and Tanks and LBs (GM2)

**Table S5. Biogas and methane production by week**

| | Biogas volume L/wk (as measured) | | | GM2 temp. | Atm. Pressure | Biogas volume L/wk (corrected to STP) | | | Methane L/wk (at STP) |
|---|---|---|---|---|---|---|---|---|---|
| Week | Total | GM1 | GM2 | °C | kPa | Total | GM1 | GM2 | Total |
| 1 | 234.8 | 72.2 | 162.6 | 23.4 | 1004.9 | 215.2 | 66.2 | 149.0 | 115.9 |
| 2 | 500.6 | 245.6 | 255.0 | 23.4 | 999.2 | 461.5 | 226.4 | 235.1 | 253.7 |
| 3 | 564.2 | 213.5 | 350.7 | 23.4 | 1003.9 | 517.6 | 195.9 | 321.8 | 280.7 |
| 4 | 578.0 | 118.8 | 459.2 | 23.4 | 1005.8 | 529.3 | 108.8 | 420.5 | 280.8 |
| 5 | 596.7 | 127.3 | 469.4 | 23.4 | 992.6 | 553.7 | 118.1 | 435.5 | 294.3 |
| 6 | 500.5 | 114.8 | 385.7 | 23.4 | 1003.5 | 459.4 | 105.4 | 354.0 | 245.6 |
| 7 | 519.8 | 104.5 | 415.3 | 23.4 | 1004.6 | 476.6 | 95.8 | 380.7 | 252.5 |

### Table S5. Biogas and methane production by week

| | | | | | | | | | |
|---|---|---|---|---|---|---|---|---|---|
| 8 | 513.6 | 74.5 | 439.1 | 23.4 | 1005.4 | 470.5 | 68.3 | 402.3 | 247.8 |
| 9 | 491.0 | 67.0 | 424.0 | 23.4 | 1004.1 | 450.4 | 61.5 | 388.9 | 237.0 |
| 10 | 526.4 | 81.0 | 445.4 | 23.4 | 1005.5 | 482.2 | 74.2 | 408.0 | 254.2 |
| 11 | 481.9 | 71.3 | 410.6 | 23.4 | 1005.1 | 441.6 | 65.3 | 376.3 | 232.4 |
| 12 | 498.9 | 73.3 | 425.6 | 23.4 | 996.1 | 461.3 | 67.8 | 393.6 | 243.1 |
| 13 | 489.5 | 80.5 | 409.0 | 23.4 | 999.8 | 450.9 | 74.2 | 376.8 | 238.0 |
| 14 | 537.7 | 83.2 | 454.5 | 23.4 | 999.5 | 495.5 | 76.7 | 418.8 | 261.3 |
| 15 | 505.4 | 65.0 | 440.4 | 23.4 | 998.9 | 466.0 | 59.9 | 406.1 | 245.0 |
| 16 | 318.3 | 23.4 | 294.7 | 23.4 | 1002.0 | 292.6 | 21.5 | 270.9 | 154.7 |
| 17 | 434.5 | 48.0 | 386.5 | 23.4 | 996.1 | 401.8 | 44.4 | 357.4 | 210.8 |
| 18 | 435.6 | 52.7 | 382.9 | 23.4 | 996.8 | 402.5 | 48.7 | 353.8 | 211.6 |
| 19 | 452.2 | 67.6 | 384.6 | 23.4 | 997.4 | 417.6 | 62.4 | 355.2 | 219.9 |
| 20 | 389.3 | 57.4 | 331.9 | 23.4 | 999.7 | 358.7 | 52.9 | 305.8 | 188.9 |
| 21 | 463.9 | 66.0 | 397.9 | 23.4 | 1001.0 | 426.9 | 60.7 | 366.1 | 243.1 |
| 22 | 441.4 | 53.4 | 388.0 | 23.4 | 1000.5 | 406.3 | 49.2 | 357.2 | 213.3 |
| 23 | 313.0 | 3.9 | 309.1 | 23.4 | 1002.3 | 287.6 | 3.6 | 284.0 | 148.9 |
| 24 | 328.6 | 37.5 | 291.1 | 23.4 | 1002.9 | 301.8 | 34.4 | 267.3 | 158.4 |
| 25 | 69.9 | 4.1 | 65.8 | 23.4 | 998.1 | 64.5 | 3.8 | 60.7 | 33.5 |
| 26 | 463.6 | 57.2 | 406.4 | 23.4 | 1004.9 | 424.9 | 52.4 | 372.5 | 223.0 |
| 27 | 542.2 | 69.9 | 472.3 | 23.4 | 1010.9 | 494.0 | 63.7 | 430.3 | 259.6 |
| 28 | 613.8 | 72.4 | 541.4 | 23.4 | 1009.6 | 560.0 | 66.0 | 493.9 | 294.0 |
| 29 | 618.7 | 75.7 | 543.0 | 23.4 | 999.6 | 570.1 | 69.7 | 500.3 | 299.1 |
| 30 | 549.5 | 42.3 | 507.2 | 23.4 | 1000.7 | 505.8 | 38.9 | 466.8 | 264.7 |
| 31 | 567.4 | 35.5 | 531.9 | 23.4 | 1007.8 | 518.5 | 32.4 | 486.1 | 270.2 |
| 32 | 471.0 | 27.6 | 443.4 | 23.4 | 999.4 | 434.1 | 25.4 | 408.6 | 225.9 |
| 33 | 415.7 | 28.2 | 387.5 | 23.4 | 1006.2 | 380.5 | 25.8 | 354.7 | 198.4 |
| 34 | 409.8 | 27.7 | 382.1 | 23.4 | 999.7 | 377.6 | 25.5 | 352.0 | 196.9 |
| 35 | 431.0 | 32.7 | 398.3 | 23.4 | 1003.6 | 395.6 | 30.0 | 365.6 | 206.3 |
| 36 | 408.8 | 26.3 | 382.5 | 23.4 | 1013.2 | 371.6 | 23.9 | 347.7 | 193.6 |
| 37 | 405.6 | 22.3 | 383.3 | 23.4 | 1008.0 | 370.6 | 20.4 | 350.2 | 193.2 |
| 38 | 352.8 | 19.2 | 333.6 | 23.4 | 997.2 | 325.9 | 17.7 | 308.1 | 169.5 |
| 39 | 335.4 | 17.7 | 317.7 | 23.4 | 998.8 | 309.3 | 16.3 | 293.0 | 161.0 |

**Table S5. Biogas and methane production by week**

| | | | | | | | | | |
|---|---|---|---|---|---|---|---|---|---|
| 40 | 296.6 | 17.8 | 278.8 | 23.4 | 1003.8 | 272.1 | 16.3 | 255.8 | 142.0 |
| 41 | 282.1 | 17.3 | 264.8 | 23.4 | 1002.7 | 259.1 | 15.9 | 243.2 | 135.0 |
| 42 | 290.2 | 16.0 | 274.2 | 23.4 | 1003.1 | 266.5 | 14.7 | 251.8 | 138.4 |
| 43 | 279.2 | 13.7 | 265.5 | 23.4 | 992.8 | 259.0 | 12.7 | 246.3 | 134.7 |
| 44 | 198.8 | 2.2 | 196.6 | 23.4 | 1006.8 | 181.9 | 2.0 | 179.9 | 94.1 |
| 45 | 167.3 | 4.1 | 163.2 | 23.4 | 993.9 | 155.0 | 3.8 | 151.2 | 80.3 |
| 46 | 140.5 | 4.7 | 135.8 | 23.4 | 1001.5 | 129.2 | 4.3 | 124.9 | 67.1 |
| 47 | 138.9 | 4.7 | 134.2 | 23.4 | 1002.4 | 127.6 | 4.3 | 123.3 | 66.4 |
| 48 | 126.3 | 5.5 | 120.8 | 23.4 | 1002.5 | 116.0 | 5.1 | 111.0 | 60.3 |
| 49 | 123.7 | 6.8 | 116.9 | 23.4 | 994.1 | 114.6 | 6.3 | 108.3 | 59.6 |
| 50 | 279.5 | 45.9 | 233.6 | 23.4 | 1004.4 | 256.3 | 42.1 | 214.2 | 135.4 |
| 51 | 347.3 | 73.1 | 274.2 | 23.4 | 1002.3 | 319.1 | 67.2 | 252.0 | 169.5 |
| 52 | 387.1 | 42.8 | 344.3 | 23.4 | 999.0 | 356.9 | 39.5 | 317.4 | 187.3 |
| 53 | 427.5 | 37.5 | 390.0 | 23.4 | 999.6 | 393.9 | 34.6 | 359.4 | 205.9 |
| 54 | 457.9 | 26.5 | 431.4 | 23.4 | 995.5 | 423.7 | 24.5 | 399.2 | 220.4 |
| 55 | 479.7 | 18.3 | 461.4 | 23.4 | 998.3 | 442.6 | 16.9 | 425.7 | 229.7 |
| 56 | 508.4 | 18.6 | 489.8 | 23.4 | 1013.6 | 462.0 | 16.9 | 445.1 | 239.9 |
| 57 | 586.5 | 19.3 | 567.1 | 23.4 | 1004.4 | 537.8 | 17.7 | 520.0 | 278.9 |
| 58 | 578.6 | 16.0 | 562.6 | 23.4 | 1002.3 | 531.7 | 14.7 | 517.0 | 276.0 |
| 59 | 561.0 | 17.3 | 543.7 | 23.4 | 995.9 | 518.8 | 16.0 | 502.8 | 269.1 |
| 60 | 546.9 | 20.4 | 526.5 | 23.4 | 1000.1 | 503.6 | 18.8 | 484.9 | 261.2 |
| 61 | 490.9 | 29.7 | 461.2 | 23.4 | 1004.8 | 450.0 | 27.2 | 422.8 | 234.4 |
| 62 | 449.1 | 34.4 | 414.7 | 23.4 | 1001.0 | 413.2 | 31.7 | 381.6 | 215.4 |
| 63 | 425.1 | 33.9 | 391.2 | 23.4 | 997.3 | 392.6 | 31.3 | 361.3 | 204.8 |
| 64 | 425.1 | 32.6 | 392.5 | 23.4 | 997.0 | 392.7 | 30.1 | 362.6 | 205.2 |
| 65 | 444.8 | 39.3 | 405.5 | 23.4 | 1001.6 | 409.0 | 36.1 | 372.9 | 213.9 |
| 66 | 421.8 | 36.4 | 385.4 | 23.4 | 1000.3 | 388.4 | 33.5 | 354.9 | 203.1 |
| 67 | 465.9 | 30.4 | 435.5 | 23.4 | 1000.7 | 428.8 | 28.0 | 400.8 | 223.5 |
| 68 | 503.5 | 36.2 | 467.3 | 23.4 | 996.7 | 465.3 | 33.5 | 431.8 | 242.8 |
| 69 | 532.6 | 43.5 | 489.1 | 23.4 | 999.5 | 490.8 | 40.1 | 450.7 | 256.4 |
| 70 | 495.4 | 38.2 | 457.2 | 23.4 | 1001.6 | 455.6 | 35.1 | 420.5 | 237.5 |
| 71 | 466.4 | 35.7 | 430.7 | 23.4 | 1001.0 | 429.1 | 32.8 | 396.3 | 224.0 |

**Table S5. Biogas and methane production by week**

| | | | | | | | | | |
|---|---|---|---|---|---|---|---|---|---|
| 72 | 494.1 | 40.0 | 454.1 | 23.4 | 1001.0 | 454.7 | 36.8 | 417.9 | 237.3 |
| 73 | 569.7 | 41.1 | 528.6 | 23.4 | 1000.6 | 524.4 | 37.8 | 486.6 | 273.5 |
| 74 | 526.2 | 51.4 | 474.8 | 23.4 | 999.4 | 484.9 | 47.4 | 437.6 | 253.7 |
| 75 | 624.5 | 53.2 | 571.3 | 23.4 | 1005.5 | 572.1 | 48.7 | 523.3 | 298.9 |
| 76 | 653.0 | 54.4 | 598.6 | 23.4 | 1006.2 | 597.8 | 49.8 | 548.0 | 312.4 |
| 77 | 634.7 | 44.8 | 589.9 | 23.4 | 1000.7 | 584.2 | 41.2 | 543.0 | 304.6 |
| 78 | 586.7 | 40.9 | 545.8 | 23.4 | 1004.2 | 538.1 | 37.5 | 500.6 | 280.5 |
| 79 | 607.3 | 48.6 | 558.7 | 23.4 | 1005.5 | 556.3 | 44.5 | 511.8 | 290.4 |
| 80 | 616.3 | 60.8 | 555.5 | 23.4 | 1004.3 | 565.2 | 55.8 | 509.5 | 296.1 |
| 81 | 686.7 | 78.0 | 608.7 | 23.4 | 1009.0 | 626.9 | 71.2 | 555.7 | 328.7 |
| 82 | 794.1 | 86.5 | 707.6 | 23.4 | 1005.1 | 727.7 | 79.3 | 648.5 | 381.1 |
| 83 | 815.7 | 82.4 | 733.5 | 23.4 | 998.4 | 752.5 | 76.0 | 676.7 | 394.2 |
| 84 | 786.2 | 93.5 | 692.7 | 23.4 | 1007.3 | 718.9 | 85.5 | 633.4 | 376.9 |
| 85 | 783.6 | 101.5 | 682.1 | 23.4 | 1008.3 | 715.8 | 92.7 | 623.1 | 376.4 |
| 86 | 798.0 | 95.1 | 702.9 | 23.4 | 1004.4 | 731.8 | 87.2 | 644.6 | 384.1 |
| 87 | 802.6 | 118.4 | 684.2 | 23.4 | 999.7 | 739.5 | 109.1 | 630.4 | 389.5 |
| 88 | 874.1 | 113.6 | 760.5 | 23.4 | 1008.4 | 798.3 | 103.8 | 694.6 | 419.3 |
| Totals | 41749 | 4584 | 37165 | | | 38363 | 4212 | 34150 | 20130 |

### Table S6. Mass balance by period

| Operating Data | | Substrates | | | Products | | | | | Mass balance % | |
|---|---|---|---|---|---|---|---|---|---|---|---|
| Period | Weeks | (a) Subs. In COD g | (b) DG out COD g | (c) Destr. COD g | (d) Conv. Cells COD g | (e) Conv. $CH_4$ COD g | (f) Total out COD g | (g) $CH_4$ out L. | (h) $CH_4$ out COD g | A f/c*100 | B (b+f)/a*100 |
| 1 | 6 - 15 | 20970 | 12714 | 8256 | 660 | 7596 | 7680 | 2457 | 7020 | 93 | 97.3 |
| 2 | 16 - 24 | 14679 | 8920 | 5759 | 461 | 5298 | 5459 | 1749 | 4999 | 94.8 | 98 |
| 3 | 25 - 31 | 14679 | 8833 | 5846 | 468 | 5379 | 5165 | 1644 | 4697 | 88.3 | 95.4 |
| 4a | 32 - 37 | 12219 | 8477 | 3742 | 299 | 3442 | 3769 | 1214 | 3470 | 100.7 | 100.2 |
| 4b | 38 - 44 | 13623 | 10561 | 3063 | 245 | 2818 | 3030 | 975 | 2785 | 98.9 | 99.8 |
| 4c | 44 - 49 | 11081 | 9558 | 1522 | 122 | 1400 | 1344 | 428 | 1222 | 88.3 | 98.4 |
| 5a | 50 - 57 | 16773 | 11088 | 5685 | 455 | 5230 | 5218 | 1667 | 4763 | 91.8 | 97.2 |
| 5b | 58 - 63 | 12580 | 8398 | 4182 | 335 | 3848 | 4509 | 1461 | 4174 | 107.8 | 102.6 |
| 5c | 64 - 70 | 12580 | 8591 | 3989 | 319 | 3670 | 4161 | 1345 | 3842 | 104.3 | 101.4 |
| 5d | 71 - 74 | 10483 | 6524 | 3959 | 317 | 3642 | 3820 | 1226 | 3503 | 96.5 | 98.7 |
| 6a | 74 - 80 | 13080 | 7637 | 5443 | 435 | 5007 | 5408 | 1740 | 4973 | 99.4 | 99.7 |
| 6b | 81 - 86 | 14079 | 7119 | 6960 | 557 | 6403 | 6961 | 2241 | 6404 | 100.0 | 100.0 |
| [a]All | 6 - 88 | 167325 | 108934 | 58391 | 4671 | 53719 | 58833 | 18957 | 54162 | 100.8±2 | 100.3±2 |
| | | | | c = a - b | d = c*0.08 | e = c*0.92 | f = d + h | | h = g/0.35l.gCOD$^{-1}$ | | |

Mass balance A = products out as % of substrate destroyed; Mass balance B = all products as % of all inputs

[a]The mass balance for the entire experiment does not equal the average of the 12 periods; minor adjustments were made to some of the inputs to the latter, to correct for a brief shutdown and weeks in which no LB was changed.

**Table S7. Substrate destruction efficiency (as VS), by substrate, vs $COD_{FW}$ addition - from coupon data**

| FW addn. | 0%$COD_{FW}$ | | | 7.9%$COD_{FW}$ | | | 12.9%$COD_{FW}$ | | | 17.2%$COD_{FW}$ | | | 21.7%$COD_{FW}$ | | | 29.3%$COD_{FW}$ | | |
|---|---|---|---|---|---|---|---|---|---|---|---|---|---|---|---|---|---|---|
| Substr. | %VS destr. | SD | n | %VS destr. | SD | n | %VS destr. | SD | n | %VS destr. | SD | n | %VS destr. | SD | n | %VS destr. | SD | n |
| CB | 13.1 | 2.3 | 5 | 26.2 | 2.6 | 6 | 45.7 | 10.3 | 6 | 36.3 | 9.7 | 33 | 47.5 | 6.8 | 6 | 49.4 | 10.7 | 3 |
| BB | 10.7 | 1.1 | 6 | 14.9 | 2.4 | 5 | 24.8 | 2.4 | 6 | 30.7 | 8.8 | 30 | 40.1 | 2.8 | 6 | 42.8 | 4.5 | 3 |
| NP | 9.7 | 1.3 | 6 | 20.1 | 3.3 | 6 | 30.0 | 2.1 | 5 | 23.7 | 12.0 | 33 | 18.7 | 2.6 | 6 | 14.4 | 2.3 | 3 |
| FP | 9.0 | 1.5 | 6 | 22.2 | 6.8 | 6 | 38.6 | 5.2 | 6 | 42.7 | 4.5 | 29 | 61.6 | 7.4 | 6 | 65.5 | 11.8 | 3 |
| BA | 4.6 | 3.3 | 2 | 3.8 | 0.9 | 2 | 8.2 | 4.6 | 2 | 7.0 | 1.9 | 10 | 6.5 | 0.4 | 2 | 7.0 | 3.1 | 1 |

**Table S8. Physical Properties of Bulking Agents BA#4 and BA#5**

| | Particle size distribution % | | | | Water retention | Bulk density |
|---|---|---|---|---|---|---|
| | >3.36mm | <3.36mm >0.50mm | <0.50mm >0.21mm | <0.21mm | gH2O/gBA | kg/L |
| BA#4 | 44.7 | 37.9 | 12.5 | 3.4 | 2.95 | 0.119 |
| BA#5 | 52.8 | 31.2 | 11.7 | 2.9 | 2.33 | 0.220 |

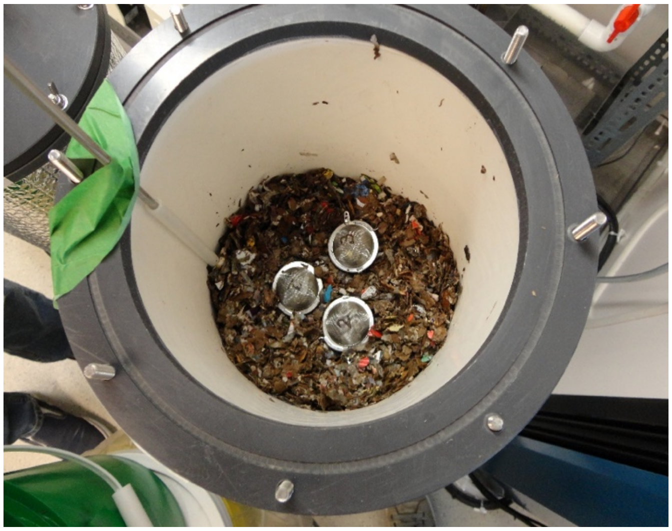

**Figure S1**: Triplicate coupons being embedded in the waste mass

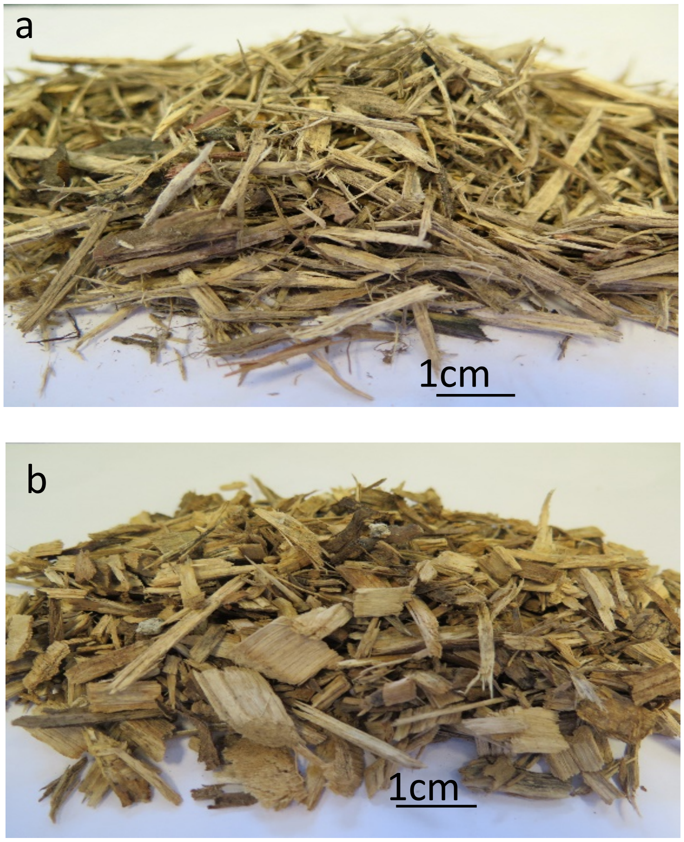

**Figure S2**: Bulking agent a) BA#4; b) BA#5; BA5 coarser and of higher bulk density

*Substrate: from elemental analysis of FW+FB+BA, average fed to Daisy (weeks 5 – 88):* $C_{90}H_{155}O_{67}N$ *(n=90 a=155 b=67 c=1), nitrogen source: ammonia.*

$d = (4n + a - 2b - 3c) = 378$

$f_e = 1 - f_s = 1 - 0.08 = 0.92$ *($f_s$ from Rittmann and McCarty for anaerobic reactions – low biomass yield)*

$r_a$: $0.125CO_2 + H^+ + e^- = 0.125CH_4 + 0.25H_2O$

$r_c$: $0.2CO_2 + 0.05NH_4^+ + 0.05HCO_3^- + H^+ + e^- = 0.05C_5H_7O_2N + 0.45H_2O$

$r_d$: $(c(n\text{-}c)/d)CO_2 + (c/d)NH_4^+ + (c/d)HCO_3^- + H^+ + e^- = (1/d)C_nH_aO_bN_c + ((2n\text{-}b+c)/d)H_2O$

$- r_d$: $0.235CO_2 + 0.00264NH_4^+ + 0.00264HCO_3^- + H^+ + e^- = 0.00264C_{90}H_{155}O_{67}N + 0.301H_2O$

$f_er_a$: $0.115CO_2 + 0.92H^+ + 0.92e^- = 0.115CH_4 + 0.23H_2O$

$f_sr_c$: $0.016CO_2 + 0.004NH_4^+ + 0.004HCO_3^- + 0.08H^+ + 0.08e^- = 0.004C_5H_7O_2N + 0.036H_2O$

$$0.00264C_{90}H_{155}O_{67}N + 0.035H_2O = 0.004C_5H_7O_2N + 0.115CH_4 + 0.104CO_2 + 0.00136NH_4^+ + 0.00136HCO_3^-$$

$$1.94C_{90}H_{155}O_{67}N + 25.7H_2O = 2.98C_5H_7O_2N + 84.6CH_4 + 76.4CO_2 + NH_4^+ + HCO_3^-$$

*COD content: substrate = 1.25 gCOD.gVS*$^{-1}$ *cells = 1.42 gCOD.gVS*$^{-1}$

*Calculated methane content: (84.6/84.6+76.4)*100 = 52.5%*

**Figure S3**. Stoichiometry of digestion of the 83-week weighted average substrate and the resultant percent methane in biogas, calculated from first principles.

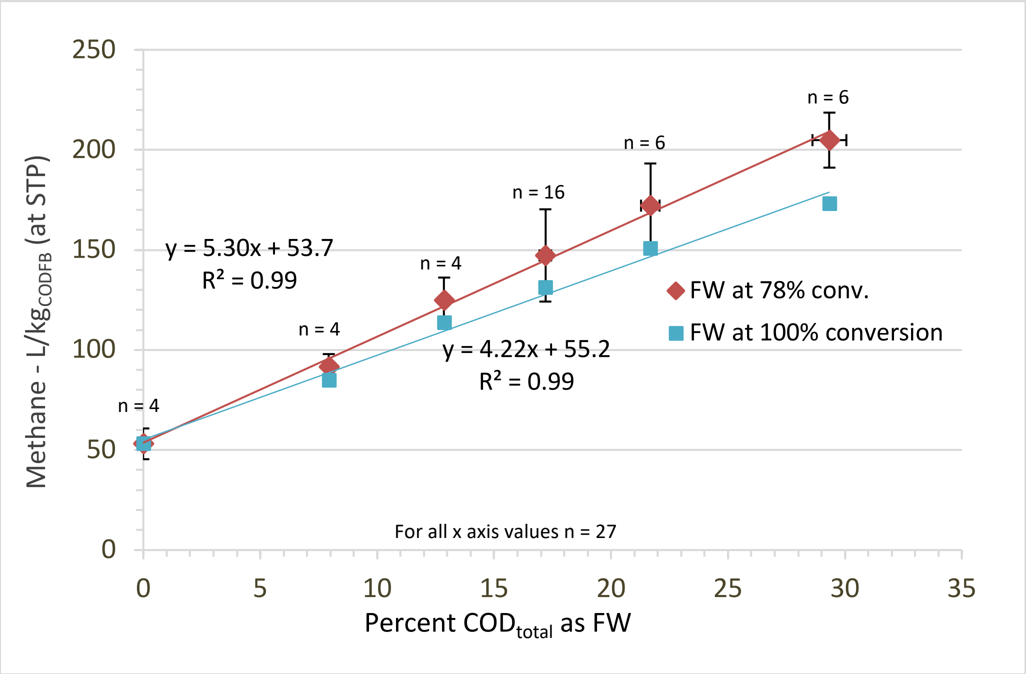

**Figure S4**: $CH_4$ produced/kg $COD_{FBadded}$ vs. percent $COD_{FWadded}$ (using Equation 11) Result at 100%$COD_{FW}$ conversion included for comparison
